# Functional role of Polymerase IV during pollen development in *Capsella*

**DOI:** 10.1101/863522

**Authors:** Zhenxing Wang, Nicolas Butel, Juan Santos-González, Filipe Borges, Jun Yi, Robert A Martienssen, German Martinez, Claudia Köhler

## Abstract

In *Arabidopsis thaliana,* the DNA-dependent RNA polymerase IV (Pol IV) is required for the formation of transposable element (TE)-derived small RNA (sRNA) transcripts. These transcripts are processed by DICER-LIKE 3 into 24-nt small interfering RNAs (siRNAs) that guide RNA-dependent DNA methylation. In the pollen grain, Pol IV is also required for the accumulation of 21/22-nt epigenetically-activated siRNAs (easiRNAs) that likely silence TEs by post-transcriptional mechanisms. Despite this proposed functional role, loss of Pol IV function in *Arabidopsis* does not cause a discernable pollen defect. Here, we show that loss of *NRPD1,* encoding the largest subunit of Pol IV in the Brassicaceae *Capsella rubella,* causes post-meiotic arrest of pollen development at the microspore stage. As in *Arabidopsis,* all TE-derived siRNAs were depleted in *Capsella nrpd1* microspores. In wild-type background, we found that the same TEs produced 21/22-nt and 24-nt siRNAs, leading us to propose that Pol IV is generating the direct precursors for 21-24-nt siRNAs, which are targeted by different DICERs. Arrest of *Capsella nrpd1* microspores was accompanied by deregulation of genes targeted by Pol IV-dependent siRNAs. The distance of TEs to genes was much closer in *Capsella rubella* compared to *Arabidopsis thaliana,* providing a possible explanation for the essential role of Pol IV for pollen development in *Capsella.* Our study in *Capsella* uncovers a functional requirement of Pol IV in microspores, emphasizing the relevance of investigating different plant models.

**One-sentence summary:** Loss of Polymerase IV function in *Capsella rubella* causes microspore arrest, revealing an important functional role of Polymerase IV during pollen development.

The author responsible for distribution of materials integral to the findings presented in this article in accordance with the policy described in the Instructions for Authors (www.plantcell.org) is: Claudia Kohler

## Introduction

In flowering plants, male and female gametes are the products of a multistep process that starts from a cell undergoing a meiotic division resulting in haploid spores dividing mitotically to form multicellular gamete-containing gametophytes. The pollen grain corresponds to the male gametophyte and forms after two mitotic divisions of the haploid microspore. The first mitotic division generates the large vegetative cell and a small generative cell that after another mitotic division will give rise to the two sperm cells. The second mitotic division can occur before pollen germination, like in *Arabidopsis thaliana,* or during pollen germination (Berger and Twell, 2011). Unlike in the male lineage where all microspores will develop into pollen, in most flowering plants only one megaspore survives and mitotically divides to give rise to the seven-celled female gametophyte containing the two female gametes, the egg cell and the central cell (Tekleyohans et al., 2017). In contrast to animals where germ cells separate early from the somatic linage, plant germ cells originate from differentiated cells that acquire the competence to undergo meiotic divisions (Schmidt et al., 2015) The formation of male and female plant gametes is connected with a partial resetting of epigenetic marks that likely serves to achieve meiotic competence (Borges and Martienssen, 2013; Baroux and Autran, 2015; Borg and Berger, 2015). Epigenetic modifications can be applied directly on the DNA in the form of DNA methylation, or on histones, the proteins that package DNA into nucleosomes. The specific type of the modification and its position on the genomic locus defines the transcriptional outcome. DNA methylation is generally (but not always) a repressive modification and is used to silence transposable elements (TEs), but also genes during specific stages of plant development. In plants, DNA methylation can occur in CG, CHG, and CHH context (where H correspond to A, T or C) and is established and maintained by different DNA methyltransferases. Methylation in CG context is maintained by the METHYLTRANSFERASE 1 (MET1), while CHG methylation maintenance requires CHROMOMETHYLASE 3 (CMT3) and to a lesser extent CMT2 (Zhang et al., 2018). The RNA-dependent DNA methylation (RdDM) pathway maintains CHH methylation by recruiting the DOMAINS REARRANGED METHYLTRANSFERASE 2 (DRM2). This pathway requires the plant-specific DNA-dependent RNA polymerases (Pol) IV and V (Herr et al., 2005; Onodera et al., 2005; Xie et al., 2004). Pol IV generates small transcripts of about 30-40-nt in size that are converted into double stranded RNA by the action of the RNA-DEPENDENT RNA POLYMERASE 2 (RDR2) (Blevins et al., 2015; Zhai et al., 2015a; Li et al., 2015). These double stranded RNAs are processed into 23- and 24-nt siRNAs by DICER-LIKE 3 (DCL3) (Xie et al., 2004; Singh et al., 2019). The 24- nt siRNAs preferentially associate with ARGONAUTE4 (AGO4) and guide DRM2 to its targets by associating with Pol V transcripts (Cao and Jacobsen, 2002; Zilberman et al., 2003; Wierzbicki et al., 2009). Recent studies further uncovered that Pol IV is required for the accumulation of 21/22-nt epigenetically activated siRNAs (easiRNAs) in pollen and establishes a hybridization barrier between plants of different ploidy grades (Martinez et al., 2018; Borges et al., 2018; Satyaki and Gehring, 2019).

Pollen formation in *Arabidopsis* is connected with reprogramming of CHH methylation. There is a strong reduction of CHH methylation during meiosis, which is followed by a restoration of CHH methylation in the pollen vegetative cell and a locus-specific restoration in sperm (Calarco et al., 2012; Ibarra et al., 2012; Walker et al., 2018). Nevertheless, CHH methylation is not completely erased during meiosis and locus­ specific CHH methylation was shown to be of functional relevance for meiosis (Walker et al., 2018). In *Arabidopsis,* the accumulation of meiosis-specific sRNAs depends on Pol IV (Huang et al., 2019) and meiotic defects have been reported in RdDM mutants *rdr2, drm2,* and *ago4,* albeit at low frequency in the Columbia (Col) accession (Oliver et al., 2016; Walker et al., 2018). There is furthermore a relaxation of heterochromatin occurring in the vegetative cell as a consequence of histone H1 depletion, which allows the DNA demethylase DEMETER (DME) to access and demethylate TEs in the vegetative cell (Slotkin et al., 2009; He et al., 2019). Demethylated TEs in the vegetative cell generate siRNAs that can move to sperm cells and may serve to enforce TE silencing in sperm (Martinez et al., 2016; Ibarra et al., 2012; Kim et al., 2019). Nevertheless, the functional relevance of enhanced TE silencing in sperm by mobile siRNAs remains to be demonstrated, since loss of DME function in pollen causes a pollen germination defect, but not a seed defect (Schoft et al., 2011).

*Arabidopsis thaliana* differs from many other species by its low repeat content of about 24% of the 135-Mb genome (Maumus and Quesneville, 2014), which may explain its apparent tolerance to the lack of Pol IV and other RdDM components. Like *Arabidopsis thaliana,* the closely related Brassicaceae *Capsella rubella* is a selfer; however, because of its recent transition to selfing (30-100 k years ago (kya)) it has maintained high numbers of TEs and almost half of the 219-Mb genome is repetitive, with many TEs being located in the vicinity to genes (Slotte et al., 2013; Foxe et al., 2009; Guo et al., 2009; Niu et al., 2019). In contrast, *Arabidopsis thaliana* became a selfer around 500 kya and experienced a strong reduction of TEs (de la Chaux et al., 2012). Given the different TE content in both species, we hypothesized that loss of Pol IV function may have a stronger impact in *Capsella rubella.* To test this hypothesis, we generated a loss-of-function allele in the *Capsella rubella* Pol IV subunit NRPD1. We found that loss of NRPD1 in *Capsella* caused impaired male fertility, as a consequence of a post-meiotic arrest of pollen at the microspore stage. Wild-type microspores accumulated Pol IV-dependent siRNAs in the size range of 21-24-nt, suggesting that the formation of easiRNAs is initiated during or after meiosis. In *Capsella* and *Arabidopsis* microspores, 21/22-nt and 24-nt siRNAs were generated from the same TE loci, suggesting that Pol IV is producing the precursors for both types of siRNAs. Consistently, we found that loss of DCL3 in *Arabidopsis* causes increased formation of 21/22-nt siRNAs, supporting the idea that different DCLs compete for the same double stranded RNA precursor molecule. Microspore arrest in *Capsella nrpd1* mutant plants correlated with a substantially stronger deregulation of genes compared to *Arabidopsis nrpd1* microspores, including known regulators of pollen development. We conclude that Pol IV in *Capsella* generates siRNAs in the size range of 21-24-nt that have important functional roles for pollen development.

## Results

### Loss of Pol IV function causes microspore arrest in *Capsella*

To test the functional role of Pol IV in *Capsella rubella,* we generated a knockout mutant in the NRPD1 subunit of Pol IV using Crispr/Cas9 (now referred to as *Cr nrpd1)* (Figure 1A). The induced deletion caused the formation of a frameshift and a stop codon after 385 amino acids, leading to a truncated protein without the catalytically active site (Onodera et al., 2005) that is likely a functional null allele. Like in the *Arabidopsis thaliana nrpd1* mutant (referred to as *At nrpd1),* TE-derived 24-nt siRNAs were abolished in *Cr nrpd1* leaves (Figure 1B) (Wierzbicki et al., 2012), connected with a strong reduction of CHH methylation levels over TEs (Figure 1C). These results reveal that Pol IV has a conserved function in siRNAs biogenesis and is required for RdDM in *Capsella* and *Arabidopsis*.

**Figure 1.**
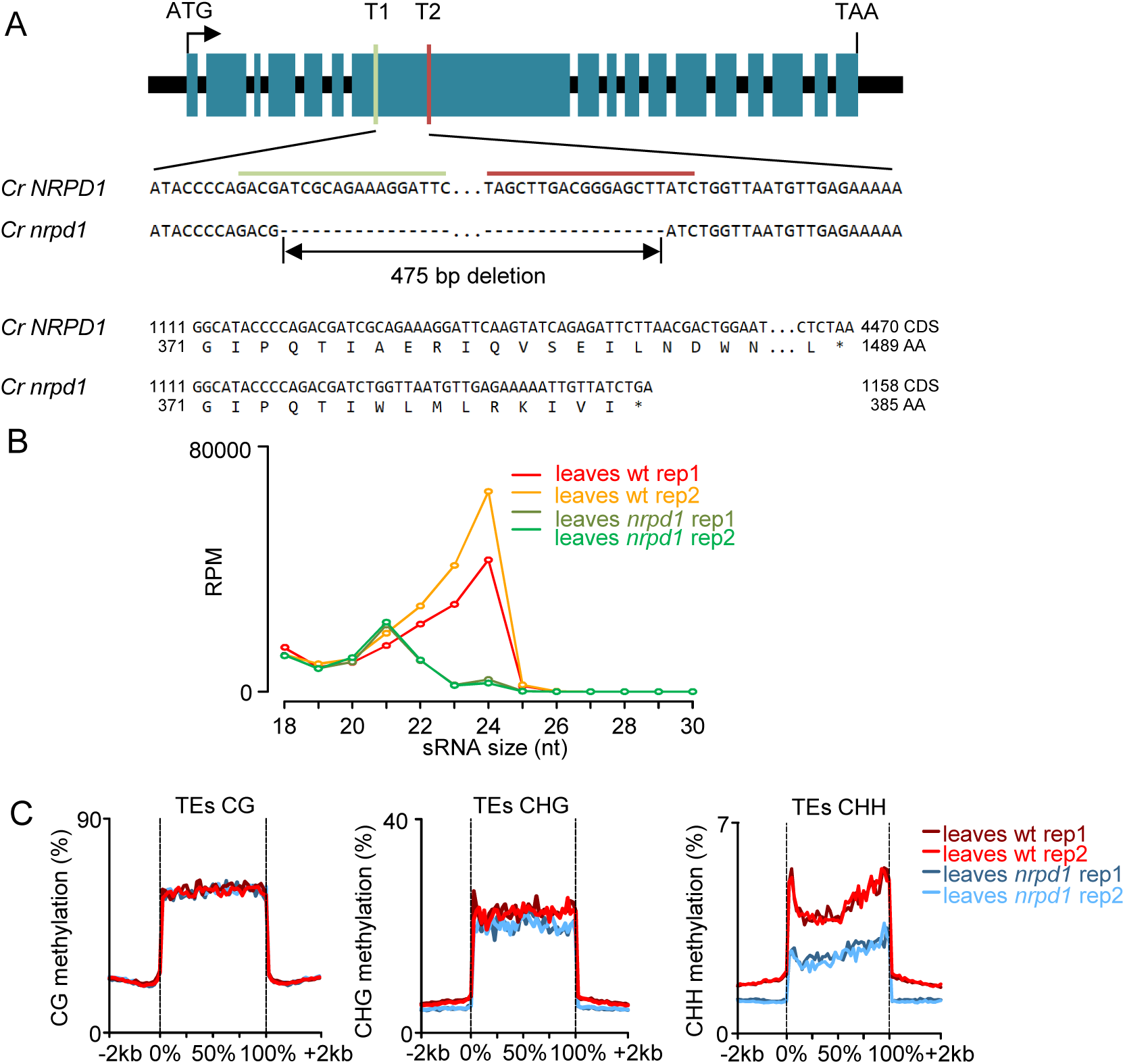
Disruption of *NRPD1* in *Capsella* impairs 24-nt siRNA formation and RdDM. (A) Deleted genomic region in *Capsella NRPD1* at 1634 - 2108 bp (genomic sequence). Target 1 (T1) and target (T2) sequences of Crispr/Cas9 are indicated. (B) Profile of TE-derived siRNAs in *Capsella* wild-type (wt) and *nrpd1* leaves. (C) DNA methylation levels at TEs in *Capsella* wt and *nrpd1* leaves.

Strikingly, homozygous *Cr nrpd1* had strongly reduced seed set (Figure 2A), on average *Cr nrpd1* siliques contained only three seeds, corresponding to about 25% of normal wild­ type seed set. We found that male fertility of *Cr nrpd1* was strongly impaired, with most pollen being arrested after meiosis at the microspore stage (Figure 3A-K). Only homozygous *Cr nrpd1* mutants were impaired in pollen development, while heterozygous *Cr nrpd1* plants were completely fertile and pollen development was normal (Figure 3C, G, K), indicating that loss of Pol IV function affects a stage prior to microspore development, or alternatively, affects tapetum development. Cross-sections of microsporangia confirmed that *Cr nrpd1* mainly formed arrested microspores, with approximate 20% of the pollen were able to complete development (Figure 3L). There were no obvious differences in tapetum development and degradation in *Cr nrpd1* compared to wild type (Figure 3M-T).

**Figure 2.**
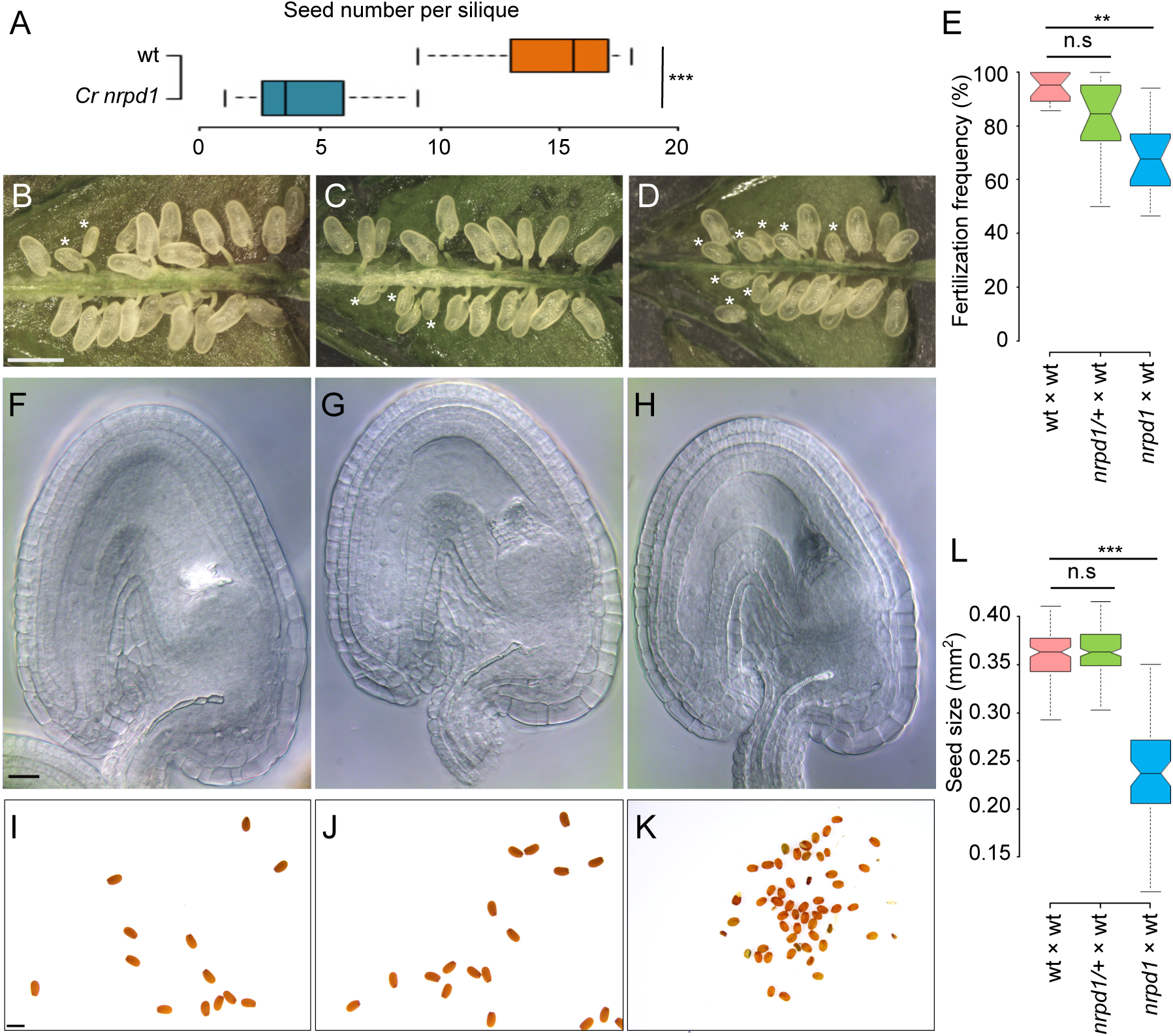
Loss of NRPD1 in *Capsella* affects female fertility and causes a reduction of seed size. (A) Total seed numbers per silique in wild-type (wt) (11 siliques) and *Cr nrpd1* mutant (24 siliques) plants. ***p <0.001 (Student’s t-test). (B-D) Siliques at 2 days after pollination (OAP) from (B) wt × wt, (C) *nrpd1*/*+* × wt, (D) *nrpd1* × wt crosses. Bar (B-D): 1mm. Asterisks mark unfertilized ovules. (E) Fertilization frequency in indicated crosses. 16 siliques per cross combination were analyzed. ** p < 0.01 (Student’s t-test). (F-H) Ovules at 2 days after emasculation of (F) wt, (G) *nrpd1*/*+,* (H) *nrpd1* plants. Bar (F-H): 20 µm. (1-K) Seeds harvested from (I) wt × wt, (J) *nrpd1*/*+* × wt, (K) *nrpd1* × wt crosses. Bar (1-K): 1mm. (L) Seed size of mature seeds derived from wt × wt (n=120), *nrpd1*/*+* × wt (n=149), *nrpd1* × wt (n=64) *** p < 0.001 (Student’s t-test). n.s, not significant.

**Figure 3.**
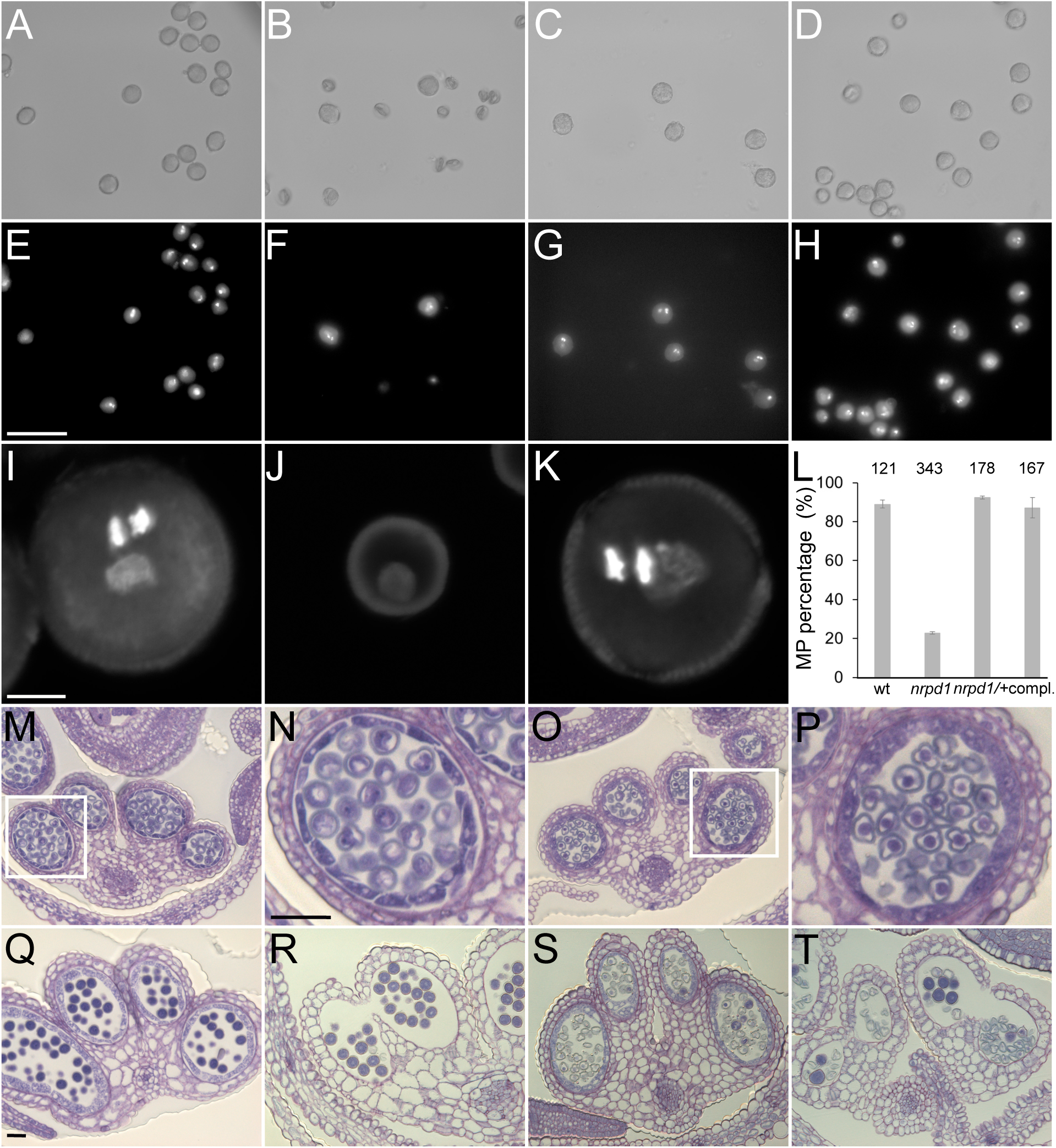
*Cr nrpd1* pollen arrest at the microspore stage. (A-D) Bright field and (E-H) corresponding DAPI staining of manually dissected pollen from anthers at stage 12/13. Pollen of wt (A and E), *Cr nrpd1* homozygotes (B and F), *Cr nrpd1* heterozygotes (C and G), and a complemented line (D and H). Bar (A-H): 50 µm. Confocal images of DAPI stained pollen of wild type (wt) (I), *Cr nrpd1* homozygotes (J), and *Cr nrpd1* heterozygotes (K). Bar (1-K): 5 µm. (L) Percentage of mature pollen (MP) in anthers dissected at stage 12/13 from wt, *Cr nrpd1* homozygotes *(nrpd1), Cr nrpd1* heterozygotes *(nrpd1*/*+)* and a complemented line (compl.). Numbers of pollen counted in each genotype were shown on top of the bars. Microsporangia cross-sections stained with Toluidine Blue at anther stage 8 (M and O), 11 (Q and S), and 12 (R and T) of wt (M, Q and R) and *Cr nrod1* (O, S and T). Bar (M,O,Q-T): 50 µm. Insets in (M) and (O) are shown enlarged in (N) (P), respectively. Bar (N and P): 50µm. wt, wild type.

Consistent with previous work showing a maternal effect of *nrpd1* mutants in *Brassica rapa* (Grover et al., 2018), about 30% of ovules of homozygous *Cr nrpd1* remained unfertilized after pollination with wild-type pollen, while there was no statistically significant fertility decrease in heterozygous *Cr nrpd1* (Figure 2B-E). Inspection of ovules of *Cr nrpd1* did not reveal obvious abnormalities; wild-type and *Cr nrpd1* ovules contained both an egg cell and unfused polar nuclei at 2 days after emasculation (Figure 2F-H). Furthermore, *Cr nrpd1* homozygous, but not heterozygous plants had strongly reduced seed size after self-fertilization or pollination with wild-type pollen (Figure 2I-L), revealing a maternal effect on ovule and seed development.

Complementation of *Cr nrpd1* with the *Arabidopsis NRPD1* genomic sequence under control of the constitutive *RPS5A* promoter (Weijers et al., 2001) fully restored pollen development in the T1 generation of transgenic plants (Figure 3D,H,L), confirming that the pollen defect is a consequence of impaired Pol IV function and that NRPD1 is functionally conserved in *Arabidopsis* and *Capsella*.

Meiotic abnormalities at low frequency were previously reported for mutants of the RdDM pathway in *Arabidopsis* (Oliver et al., 2016; Walker et al., 2018). However, we did not detect abnormal chromosome segregation during male meiosis in *Cr nrpd1* (Supplemental Figure 1) and inspection of meiotic products revealed the formation of tetrads (Supplemental Figure 2), indicating that the pollen arrest after meiosis is not a consequence of a chromosome segregation defect.

### Pol IV-dependent silencing of TEs in *Capsella* microspores

The arrest of *Cr nrpd1* pollen at the microspore stage prompted us to compare sRNAs of wild-type and *Cr nrpd1* microspores. We enriched for microspores using a percoll gradient following previously established procedures (Dupl’akova et al., 2016). On average, the purity of the fractions was 84% (Supplemental Figure 3A). Sequencing of isolated sRNAs revealed that TE-derived siRNAs in the size range of 21-24-nt were abolished in *Cr nrpd1* microspores (Figure 4A, Supplemental Figure 4A). Like previously described for *Arabidopsis* meiocytes (Huang et al., 2019), we observed a strong accumulation of 23-nt siRNAs in *Capsella* microspores (Figure 4A, Figure 5A). Nearly all TE loci generating 21/22-nt siRNAs also formed 24-nt siRNAs (Figure 4B). To test the functional role of Pol IV-dependent siRNAs in TE silencing, we isolated RNA from wild-type and mutant microspores and performed an RNA-seq analysis. Those TEs that lost 21/22- and 24-nt siRNAs, had higher transcript levels in *Cr nrpd1* microspores compared to wild type (Figure 4C), revealing a role of Pol IV-dependent siRNAs in TE silencing in microspores. Together, we conclude that Pol IV-dependent TE-derived siRNAs in the size range of 21- 24-nt are present in microspores and are required for TE silencing. This is consistent with recent work proposing that pollen easiRNAs are produced during or shortly after meiosis (Borges et al., 2018).

**Figure 4.**
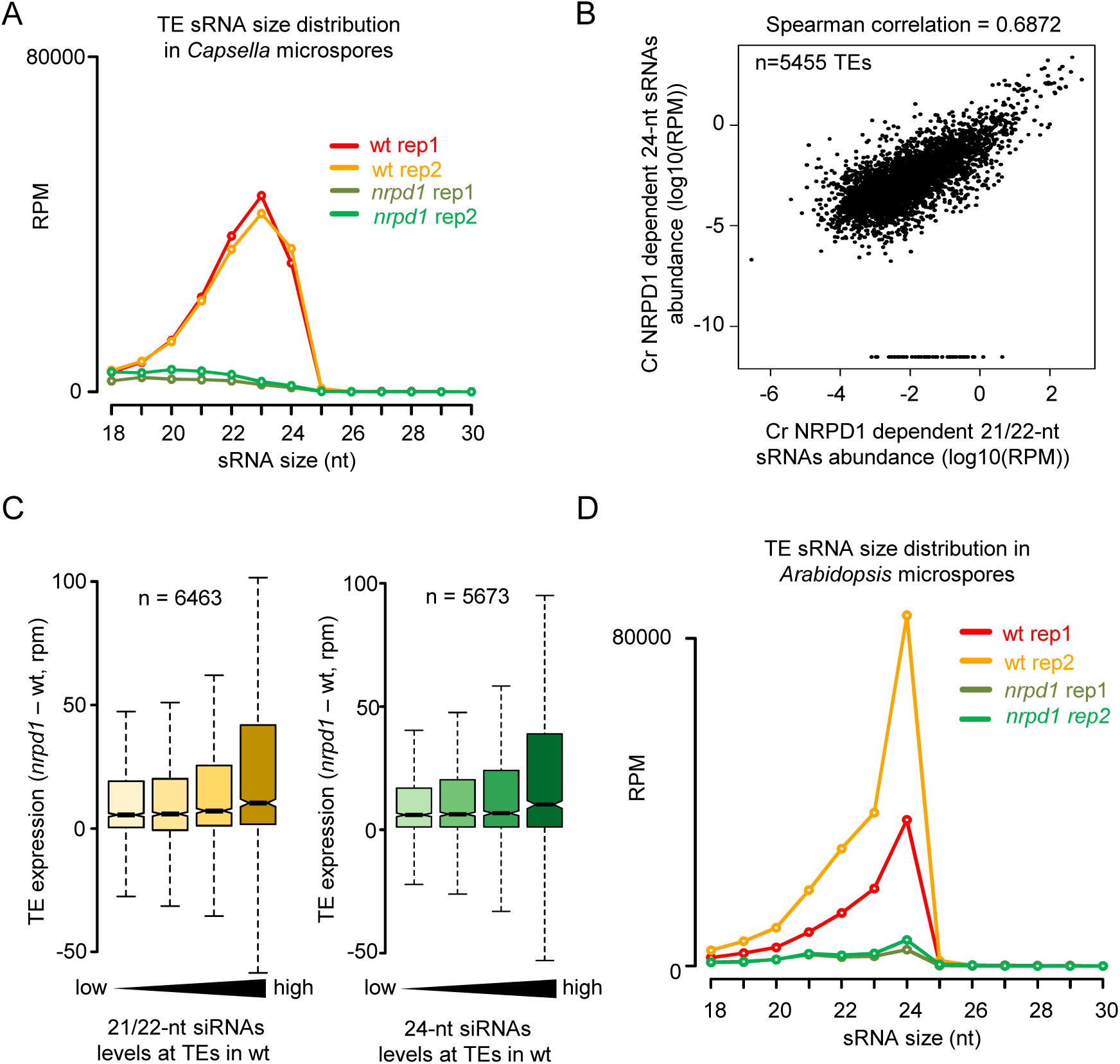
Pol IV is required for 21-24-nt siRNAs in *Capsella* microspores. (A) Profile of TE- derived siRNAs in *Capsella* wt and *nrpd1* microspores. (B) Abundance of TE-derived Cr NRPD1-dependent 21-22-nt siRNAs and 24-nt siRNAs in *Capsella* microspores. Values are indicated as the log10 of the average RPM of both libraries. Each dot represents one TE for a total of 5455 TEs. The correlation has been tested by a Spearman test (correlation coefficient 0.6872). (C) Loss of 21/22-nt and 24-nt siRNAs at TEs associates with increased transcript level of TEs in *Cr nrpd1* microspores. Increasing accumulation of siRNAs over TEs is plotted from low to high levels of accumulation. Only TEs with siRNAs more in wt than in *Cr nrpd1* are represented. Differences between first and last categories are significant *(P* = 3.4e-13 and 1.4e-9, respectively, Wilcoxon test). (D) sRNA profile of TE-derived siRNAs from *Arabidopsis* wt and *nrpd1* microspores.

**Figure 5.**
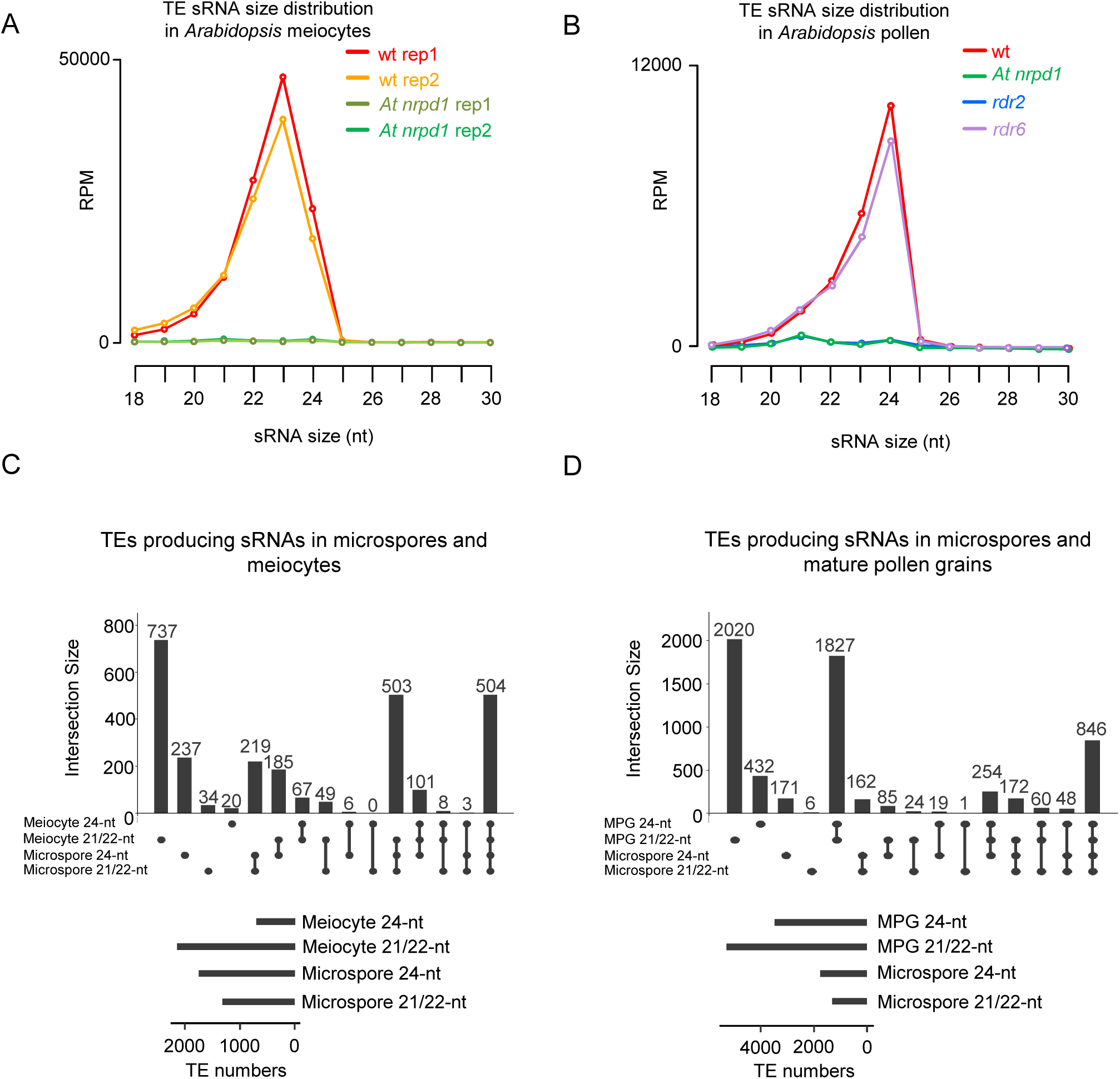
Meiocytes, microspores and mature pollen grain accumulate overlapping sets of siRNAs. (A) TE-derived siRNA distribution in *Arabidopsis* meiocytes of the indicated genetic background (data from Huang et al., 2019). (B) TE-derived siRNA distribution in *Arabidopsis* pollen grains of the indicated genetic background. (C) Upset plot showing the overlap of TEs accumulating 21/22-nt siRNAs or 24-nt siRNAs in *Arabidopsis* microspores and meiocytes (data from Huang et al., 2019). (D) Upset plot showing the overlap of TEs accumulating 21/22-nt siRNAs or 24-nt siRNAs in *Arabidopsis* microspores and mature pollen grain (MPG) (data from Martinez et al., 2018).

### Pol IV-dependent siRNAs also accumulate in *Arabidopsis* microspores

Previous work demonstrated that Pol IV generates small transcripts in the size range of 30-40-nt that are converted into 24-nt siRNAs by the action of DCL3 (Zhai et al., 2015a; Blevins et al., 2015; Yang et al., 2016). To be able to genetically dissect the Pol IV-dependent pathway leading to the formation of siRNAs in the size range of 21-24-nt, we tested for the presence of similar siRNAs in *Arabidopsis* microspores.

We sequenced sRNAs from *Arabidopsis* wild-type and *nrpd1* microspores that had been enriched to 90% following the same procedures as applied for *Capsella* (Supplemental Figure 3B, Supplemental Figure 4B). Like in *Capsella* microspores, *Arabidopsis* microspores accumulated TE-derived siRNAs in the size range of 21-24-nt that were abolished in *At nrpd1* microspores (Figure 4D). Thus, 21/22-nt TE-derived siRNAs are already present in microspores and depend on Pol IV, strongly supporting the idea that biogenesis of easiRNAs present in mature pollen starts at an earlier stage, most likely during meiosis. Consistently, microspores and meiocytes as well as microspores and mature pollen grains share a large number of loci generating Pol IV-dependent siRNAs (Figure 5C, D).

To test the functional requirement of Pol IV-derived siRNAs in TE silencing, we correlated the TEs producing Pol IV-dependent 21/22-nt and 24-nt siRNAs to TE expression changes in *At nrpd1,* lacking Pol IV function. We found a significant association between Pol IV-dependent siRNAs and expression change of TEs in *At nrpd1* microspores (Figure 6A) similar to *Capsella* (Figure 4C), revealing that TE-derived siRNAs are involved in the repression of TE expression in microspores.

**Figure 6.**
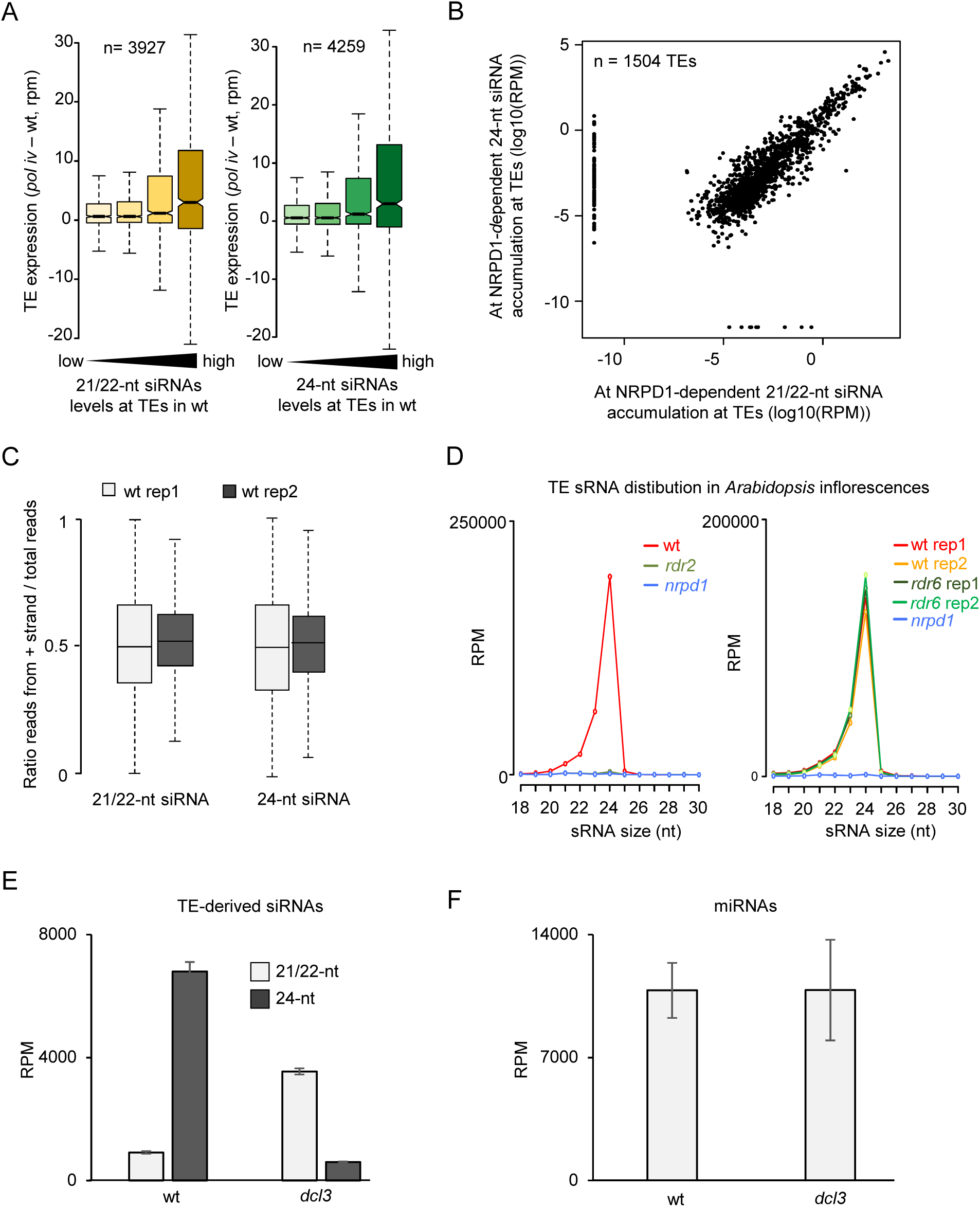
Pol lV/RDR2 generate templates for 21-24-nt siRNAs. (A) Loss of Pol IV-dependent 21/22-nt easiRNAs associates with increased transcript levels of TEs in *Arabidopsis* microspores. Increasing accumulation of siRNAs over TEs is plotted from low to high levels of accumulation. In both plots, siRNAs levels at TEs in wt increase from left to right in quantiles. Differences between first and last categories are significant (p = 2.6e-10 and 1.5 e-14, respectively, Wilcoxon test). (B) Abundance of At NRPD1-dependent 21-22-nt siRNAs and 24-nt siRNAs at TEs in *Arabidopsis* microspores. Values are indicated as log10 of the average reads per million (RPM) of both libraries. Each dot represents one TE for a total of 1504 TEs. The correlation has been tested by a Spearman test (correlation coefficient 0.7686). (C) Plots showing the distribution of the ratio of the number of reads mapped against the positive strand to the total number of mapped reads. Left plots shows analysis for 21/22-nt reads, right plot for 24-nt reads. (D) TE-derived siRNA distribution in inflorescences of *rdr2* (left panel; data from Zhai et al., 2015) and *rdr6* (right panel; data from Panda et al., 2016). (E) Average total 21/22-nt or 24-nt reads mapping against TEs in wt or *dcl3* libraries (data from Li et al., 2015). Reads were normalized to show RPM values. (F) Average total 21/22-nt reads mapping against miRNAs in wt or *dcl3* libraries (data from Li et al., 2015). Reads were normalized to show RPM values.

Pol IV is usually associated with 24-nt siRNAs through the RdDM pathway and its strong effect on the production of 21/22-nt siRNAs in pollen is thus unexpected. One hypothesis could be that Pol IV transcripts are direct precursors of 21/22-nt siRNAs. If true, we expected that microspore TE-derived 21/22-nt and 24-nt siRNAs should arise from the same genomic loci. Consistently, like in *Capsella* microspores (Figure 4B), nearly all TE loci generating 21/22-nt siRNAs also formed 24-nt siRNAs (Figure 6B). Visualizing the individual reads in a genome browser showed that all read sizes accumulated along the same loci (Supplemental Figure 5). Taken together, this data show that TE-derived 21/22- nt siRNAs and 24-nt siRNAs are produced from the same loci in a Pol IV-dependent manner and are able to repress TEs in both *Capsella* and *Arabidopsis*.

### TE-derived siRNAs in microspores require RDR2 activity

TE-derived siRNAs were produced from both DNA strands (Figure 6C and Supplemental Figure 5), suggesting the involvement of an RNA-dependent RNA polymerase in the production of the double stranded RNAs used as template for the microspore TE-derived siRNA production.

Among the three RDRs with known function in *Arabidopsis,* RDR2 is tightly associated with Pol IV (Li et al., 2015; Zhai et al., 2015a) and RDR6 has been shown to affect some easiRNAs (Creasey et al., 2014; Martinez et al., 2018). To assess the potential involvement of RDR2 and RDR6 in TE-derived siRNA production in microspores, we analyzed publically available sRNA sequencing data from *rdr2* and *rdr6* inflorescences (Zhai et al., 2015a; Panda et al., 2016). The sRNA pattern of wild-type and *At nrpd1* inflorescence tissues was comparable to that of microspores (Figure 4D and Figure 6D), indicating that the siRNAs identified in *rdr2* and *rdr6* inflorescences are comparable to those of microspores. While *rdr6* had no effect on the distribution of TE-derived siRNAs, in *rdr2* inflorescences the accumulation of TE-derived siRNAs was abolished (Figure 6D), indicating that RDR2 is likely involved in TE siRNA biogenesis. We also generated siRNA profiles from *rdr2* and *rdr6* pollen (Figure 5B), which confirm the data obtained from inflorescences and reveal that RDR2, but not RDR6 is required for the generation of 21/22-nt siRNAs. These results reinforce the idea that 21/22-nt and 24-nt TE-derived siRNAs present in pollen are processed from a double stranded RNA produced by Pol IV and RDR2.

### DICERs producing TE-derived siRNAs compete for the same double stranded RNA template

Our data suggests that TE-derived siRNAs of different size classes are derived from double stranded RNAs produced by Pol IV/RDR2 (Figure 6B). We hypothesized that the production of different size classes of siRNAs is a consequence of different DICERs competing for the same double stranded RNA template, as it has been previously shown to occur upon disruption of DCL3 function (Gasciolli et al., 2005; Henderson et al., 2006; Kasschau et al., 2007; Bond and Baulcombe, 2015). If true, the impairment of DCL3 should increase the proportion of Pol IV-dependent 21/22nt siRNAs accumulating over defined loci.

To test this idea, we used publically available sRNA sequencing data of *dcl3* inflorescences (Li et al., 2015). We quantified the number of normalized 21/22-nt siRNAs and 24-nt siRNAs mapped against TEs in wild-type and *dcl3* inflorescences. In wild type, 24-nt siRNAs were the most abundant siRNA, exceeding the level of 21/22-nt siRNAs by nearly seven fold (Figure 6E). In *dcl3,* the abundance of 21/22-nt siRNAs was highly increased, while 24-nt siRNAs were depleted (Figure 6E). To rule out that the increased abundance of 21/22-nt siRNAs is a consequence of a normalization artifact, we performed the same analysis with miRNAs. As DCL3 is not involved in the 21/22-nt miRNA pathway, observed changes would be suggestive for a normalization artifact. The abundance of 21/22-nt miRNAs was highly similar in wild type and *dcl3* (Figure 6F), strongly supporting the notion that the observed increase of 21/22-nt siRNAs in *dcl3* inflorescences is not a consequence of a normalization problem. These results indicate that there is indeed a competition between DCL3 and other DCLs for the same double stranded RNA precursor, in agreement with previous data (Gasciolli et al., 2005; Henderson et al., 2006; Kasschau et al., 2007; Bond and Baulcombe, 2015).

### TE-derived siRNAs are highly enriched at *COPIA95* in *Capsella* microspores

To investigate whether Pol IV-dependent siRNAs target similar loci in *Arabidopsis* and *Capsella,* we identified sRNA reads mapping to both genomes, which were then mapped to 527 *Arabidopsis* TE consensus sequences previously reported (Repbase) (Bao et al., 2015). We found a similar number of TE families accumulating Pol IV-dependent siRNAs in *Arabidopsis* and *Capsella* microspores, (316 and 301 TE families, respectively), of which 225 were common between both species (Figure 7A). There were substantially fewer TE families forming 24-nt siRNAs in *Capsella* microspores (133) compared to *Arabidopsis* (303), but the majority of those overlapped between both species (103). Nearly all TE families forming 24-nt siRNAs also formed 21/22-nt siRNAs (94.1% (285 out of 303 TE families) in *Arabidopsis* and 99.2% (132 out of 133 TE families) in *Capsella)* (Figure 7A), supporting the idea that 21/22-nt siRNAs and 24-nt siRNAs are derived from the same TE loci in microspores.

**Figure 7.**
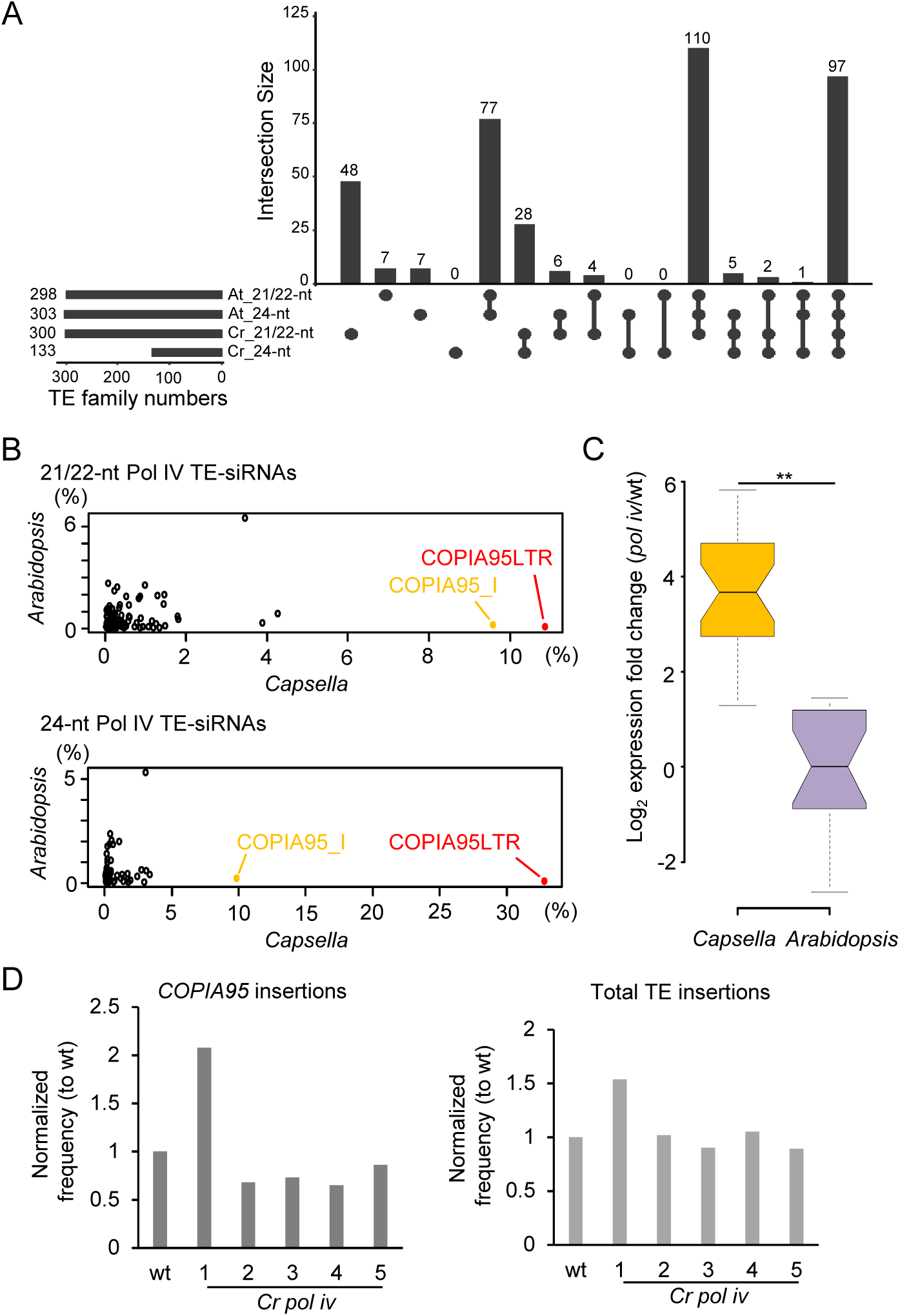
COP/A95-siRNAs are highly enriched in *Capsella* microspores. (A) Upset plots of TE families accumulating Pol IV-dependent 21/22-nt and 24-nt siRNAs in *Arabidopsis (At)* and *Capsella (Cr).* (B) Proportions of Pol IV-dependent 21/22-nt and 24-nt siRNAs accumulating at specific TE consensus sequences in relation to all TE­ siRNAs. Reads mapping to *COPIA95* long-terminal repeats (LTR) and internal (I) sequences are highlighted in red and yellow, respectively. (C) Log2 expression fold change of mRNAs for *COPIA95* elements in *nrpd1* mutant microspores of *Arabidopsis* and *Capsella* compared to the corresponding wild type. ** p < 0.01 (Student’s t-test). (D) Relative number of *COPIA95* insertions (left panel) and total TE insertions (right panel) compared to the corresponding wild-type control in five progenies of homozygous *Cr nrpd1*.

To investigate the specificity of TE-derived siRNAs in *Arabidopsis* and *Capsella* microspores, we calculated the proportion of siRNAs targeting specific TE families. Strikingly, we found that nearly 20% of 21/22-nt siRNAs and more than 40% of 24-nt siRNAs were derived from the *COPIA95* family *(COPIA95* long-terminal repeats (LTR) and internal (I)) in *Capsella* microspores, while only 0.3% of both siRNA classes were derived from *COPIA95* in *Arabidopsis* microspores (Figure 7B). We identified 17 and 70 TEs accumulating COPIA95-derived Pol IV-dependent siRNAs in *Arabidopsis* and *Capsella* respectively, indicating an expansion of the *COPIA95* TE family in *Capsella* (Supplemental dataset 1). The prominent targeting of *COPIA95* in *Capsella* microspores by Pol IV-dependent siRNAs made us address the question whether loss of Pol IV function may cause increased expression and transposition of *COPIA95.* Indeed, *COPIA95* was highly upregulated in *Cr nrpd1* microspores, but remained silenced in microspores of *At nrpd1* (Figure 7C). To test whether increased expression caused heritable transposition, we performed whole genome sequencing of five homozygous *Cr nrpd1* mutants that were derived from homozygous *Cr nrpd1* parental plants. We mapped genomic reads to *COPIA95* elements and found that one of the five tested mutants had a two-fold increase of *COPIA95* elements (Figure 7D). We corroborated this result by using TEPID to identify *de novo* insertions (Stuart el al 2016) and also detected increased insertions in the same mutant plant (Figure 7D). This result supports the idea that Pol IV is required to prevent TE remobilization in *Capsella* and is, in particular, required to silence *COPIA95*.

### Loss of Pol IV causes transcriptional changes in microspores

To understand the cause for post-meiotic arrest of *Capsella* microspores, we compared the transcriptome changes in *At* and *Cr nrpd1* microspores. We found a comparable number of genes being upregulated (log2 fold-change >1, p<0.05) in *nrpd1* microspores of both species; there were however about twice as many genes downregulated (log2 fold-change <-1, p<0.05) in *Cr nrpd1* microspores compared to *At nrpd1* (Supplemental Figure 6A, B; Supplemental dataset 2). While there was no significant overlap between downregulated genes in *Cr* and *At nrpd1* microspores (Supplemental Figure 6B), there was a significant overlap of upregulated genes in *Cr* and *At nrpd1,* with a significant enrichment for genes with functional roles in stimulus response, cell wall organization, and defense responses (Supplemental Figure 6C).

We tested whether deregulated genes in *Cr* and *At nrpd1* microspores were targeted by 21/22-nt or 24-nt siRNAs. We found a significant overlap between upregulated genes and downregulated genes with genes losing 21/22-nt and 24-nt siRNAs in *Cr nrpd1* (Figure 8A, Supplemental dataset 3). In contrast, in *At nrpd1* only downregulated genes significantly overlapped with genes losing 21/22-nt and 24-nt siRNAs (Supplemental Figure 6D, Supplemental dataset 4). Upregulated genes losing 21/22-nt or 24-nt siRNAs in *Cr nrpd1* had functional roles in proteolysis and catabolic processes, cell killing, and interspecies organismal interactions (Figure 8B). The distance of TEs to neighboring genes was significantly shorter in *Capsella* compared to *Arabidopsis,* independently of their direction of deregulation (Figure 8C; Wilcoxon test, p<2e-15). Nevertheless, upregulated genes in *Capsella* had an even shorter distance to neighboring TE than non­ deregulated or downregulated genes, suggesting an impact of neighboring TEs on gene expression in *Cr nrpd1* microspores. There was however no preference for *COPIA95* among those TEs being close to deregulated genes (p=1, hypergeometric test).

**Figure 8.**
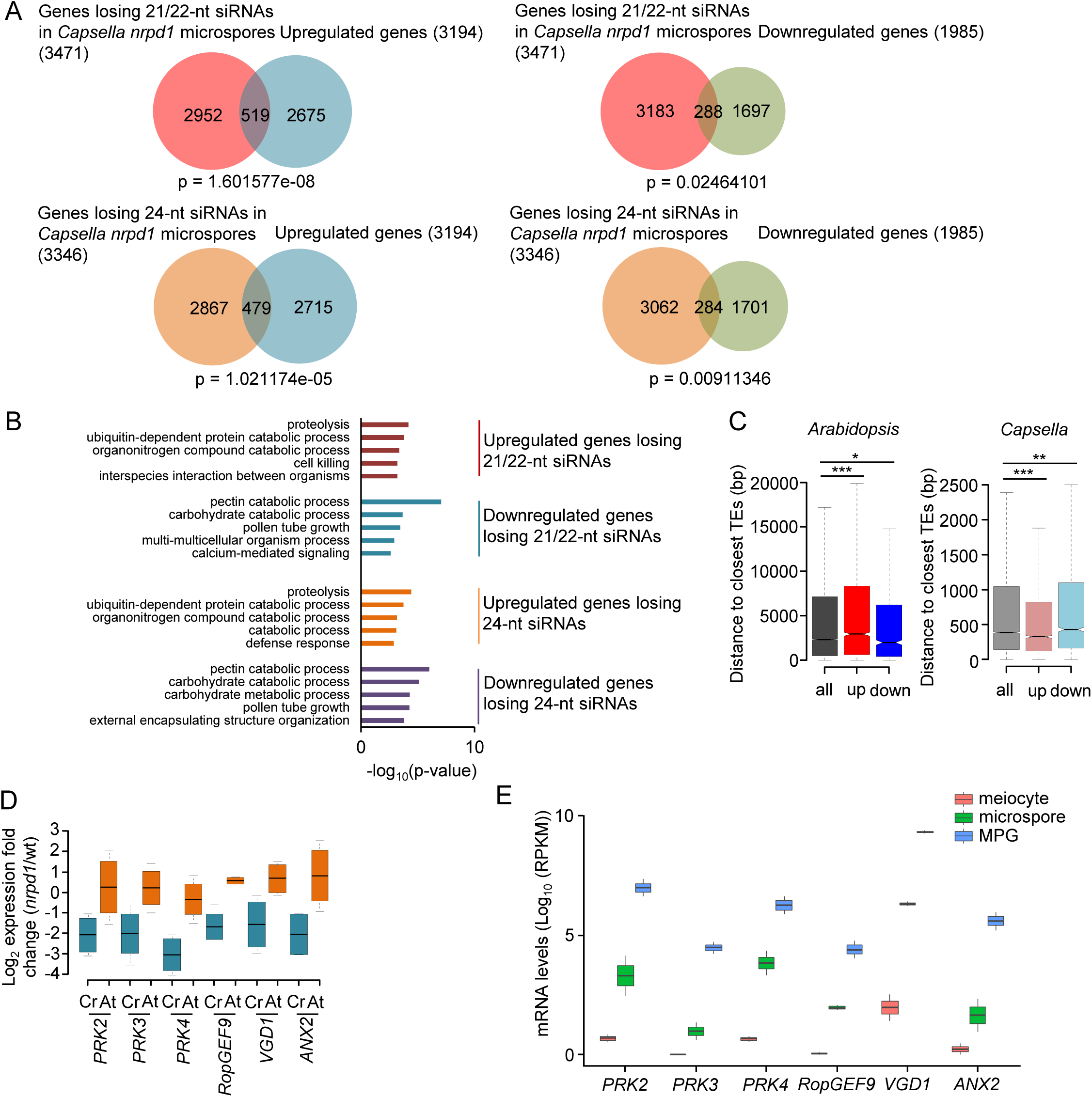
Deregulated genes differ in *Arabidopsis* and *Capsella nrpd1* mutant microspores. (A) Venn diagrams showing overlap of deregulated genes (|log_2_ fold change|> 1, p < 0.05) in *nrpd1* microspores of *Capsella* and genes losing 21/22-nt and 24-nt siRNAs at 2kb up­ and downstream and gene body (log_2_ fold change < −1, p < 0.05) in *Capsella nrpd1* microspores. (B) Enriched gene ontologies (GOs) for biological processes of intersected genes losing siRNAs and deregulated genes in *Capsella nrpd1* microspores. Top 5 GOs of each analysis are shown. (C) Distance of *Arabidopsis* and *Capsella* genes to closest TEs. All: all genes, up: significantly upregulated genes, down: significantly downregulated genes. * p < 0.05, ** p < 0.01, *** p < 0.001, n.s, not significant. (Statistical analysis: Wilcoxon test). (D) Log_2_ expression fold change of *VGD1, PRK2, PRK3, CHX21, JGB* and *ANX2* genes in *nrpd1* microspores compared to wild type (wt) in *Capsella (Cr)* and *Arabidopsis (At).* (E) mRNA levels of *VGD1, PRK2, PRK3, CHX21, JGB* and *ANX2* in *Arabidopsis* wild-type meiocytes, microspores and mature pollen grain (MPG). We added plus 1 to all values to avoid negative log_10_ values.

Interestingly, we found that downregulated genes associated with loss of 21/22-nt and 24-nt siRNAs in *Cr nrpd1* were enriched for genes involved in pollination; among those were known regulators of pollen tube growth like VANGUARD1 (VGD1), ANXUR2 (ANX2), CATION/H+ EXCHANGER 21 (CHX21), and JINGUBANG (JGB) (Figure 8D, Supplemental Figure 7; Jiang et al., 2005; Boisson-Dernier et al., 2005; Lu et al., 2011; Ju et al., 2016). We furthermore identified homologs of pollen receptor kinase encoding genes *PRK2* and *PRK3* among downregulated genes losing 21/22-nt and 24-nt siRNAs (Figure 8D, Supplemental Figure 7). While there was also a significant overlap of downregulated genes in *At nrpd1* with genes losing siRNAs (Supplemental Figure 6D), those genes were not enriched for genes involved in pollination (Supplemental Figure 6E) and the aforementioned genes were not deregulated in *At nrpd1* (Figure 8D). The affected pollen receptor kinases have partly redundant functions in pollen tube growth and perception of female attractant peptides (Chang et al., 2013; Takeuchi and Higashiyama, 2016). Importantly, RNAi-mediated knockdown of *PiPRK1,* a PRK homologue Petunia, causes microspore arrest (Lee et al., 1996), suggesting that reduced expression of PRKs may contribute to the *Cr nrpd1* microspore arrest. All genes were highly induced in the microspores to mature pollen transition (Figure 8E), suggesting that their expression is required to ensure viable pollen formation, a hypothesis that remains to be tested.

## Discussion

In this manuscript, we report that loss of Pol IV function in *Capsella rubella* causes arrest of microspore development and a maternal effect on ovule and seed development, strongly differing from the lack of obvious reproductive abnormalities of *nrpd1* mutants in *Arabidopsis* (Mosher et al., 2009). Previous work revealed that mutations in NRPD1, NRPE1 and RDR2 in *Brassica rapa* cause a maternal effect on seed development, while no defect in pollen development was reported (Grover et al., 2018). The mutation in *B. rapa NRPD1* was not a null allele; however, the mutation in *RDR2* completely abolished production of 24-nt siRNAs (Grover et al., 2018), indicating that this mutant was a functional null for *RDR2.* Since loss of *Cr NRPD1* caused a similar molecular effect as mutations in *Arabidopsis* and *B. rapa NRPD1* (depletion of 24-nt siRNAs and CHH methylation) (Wierzbicki et al., 2012; Panda et al., 2016; Grover et al., 2018) and that the *Cr nrpd1* mutant could be complemented with the *Arabidopsis NRPD1* sequence, we conclude that the molecular function of Pol IV is likely conserved between the three species, but the targets differ. Interestingly, loss of Pol IV function in tomato also causes sterility, but the cause for this phenotype remains to be explored (Gouil and Baulcombe, 2016). The microspore arrest in *Cr nrpd1* is possibly a consequence of TEs being in close vicinity to either essential regulators of microspore development, like PRKs, or genes that cause microspore arrest upon overexpression. The distance of TEs to neighboring genes is substantially larger in *Arabidopsis* compared to *Capsella,* supporting this notion.

We observed a heritable remobilization of the *COPIA95* element in progenies of *Cr nrpd1* mutant, consistent with this element being preferentially targeted by Pol IV-generated siRNAs in *Capsella* and strongly activated in *Cr nrpd1* microspores. Interestingly, in *Arabidopsis,* the *COPIA* element *ONSEN* also undergoes transgenerational retrotransposition in *nrpd1* after heat treatment and new *ONSEN* insertions differ between siblings derived from a single plant (Ito et al., 2011). Since amplification of *COPIA 95* was only observed in one of the five tested *Cr nrpd1* progenies, it seems unlikely that the consistently observed microspore arrest is connected to TE remobilization. Alternatively, it is possible that only those microspores survive where TE remobilization did not occur, or did occur at low frequency.

*Cr nrpd1* microspores were completely depleted for TE-derived siRNAs, including 21/22- nt siRNAs, which are usually not associated with Pol IV (Xie et al., 2004; Zhai et al., 2015a; Blevins et al., 2015). A similar depletion of 21/22-nt siRNAs (easiRNAs) was previously reported in the mature pollen grain of *At nrpd1* mutants (Martinez et al., 2018; Borges et al., 2018). The biogenesis of easiRNAs was suggested to be a consequence of reduced heterochromatin formation in the vegetative cell and resulting TE activation (Slotkin et al., 2009; Creasey et al., 2014). Based on genetic data Borges et al. (2018) proposed that easiRNA biogenesis occurs earlier, during or early after meiosis. Our data reveal that 21/22-nt TE-derived siRNAs are already present in the microspores and given the similarity to meiocyte siRNAs (Huang et al., 2019), they are likely generated before or during meiosis.

In rice and maize, highly abundant 21-nt phasiRNAs accumulate in premeiotic anthers and 24-nt phasiRNAs are enriched in meiotic-stage anthers (Zhai et al., 2015b; Johnson et al., 2009; Komiya et al., 2014). The 21-nt phasiRNAs were shown to be important for male fertility in rice and disruption of 24-nt phasiRNA production yields conditional male sterility in maize (Fan et al., 2016; Teng et al., 2018). Biogenesis of premeiotic and meiotic phasiRNAs in maize seems to occur in the tapetum, rather than in meiocytes where they accumulate (Zhai et al., 2015b). The production of 24-nt phasiRNAs depends on *miR2275,* a pathway that is widely present in the eudicots, but missing in the *Brassicaceae* (Xia et al., 2019), suggesting evolutionary divergence of the functional role of phasiRNAs during pollen development. Cross-sections did not reveal obvious tapetal defects in *Cr nrpd1,* indicating that microspore arrest in *Cr nrpd1* is not a consequence of a tapetal defect.

The strong dependency of TE-derived siRNA accumulation on Pol IV suggests that Pol IV transcripts are the precursors for all sizes of TE-derived siRNAs in microspores. Previous work revealed that in the absence of DCL3, other DCL proteins (DCL1, DCL2, and DCL4) are able to process Pol IV transcripts into 21- or 22-nt siRNAs (Gasciolli et al., 2005; Henderson et al., 2006; Kasschau et al., 2007; Bond and Baulcombe, 2015). We thus propose that before or during meiosis, Pol IV transcripts are targeted by other DCLs than only DCL3, explaining why all sizes of Pol IV-dependent siRNAs derive from the same TE loci.

In *Arabidopsis* siliques, a nuclear localized form of DCL4 was shown to target Pol IV transcripts and generates 21-nt siRNAs (Pumplin et al., 2016). The abundance of those 21-nt Pol IV-derived siRNAs was nevertheless low, contrasting to the high abundance in microspores. One possible explanation could be that the disruption of the nuclear envelope during meiosis allows cytoplasmic DCLs to gain access to Pol IV/RDR2 transcripts. This implicates that meiosis is the trigger of Pol IV-dependent 21-24-nt siRNA production, consistent with our genetic data. Not mutually exclusive with this scenario is the possibility that 22-nt siRNAs produced during meiosis trigger secondary 21/22-nt siRNA production in the mature pollen grain by targeting TEs transcripts expressed in the vegetative cell of pollen (Slotkin et al., 2009). This amplification of the signal by the canonical post-transcriptional gene silencing (PTGS) pathway (Martinez de Alba et al., 2013) should result in high abundant 21/22-nt siRNAs in mature pollen, which is in agreement with published siRNA profiles of pollen (Martinez et al., 2018; Borges et al., 2018).

We have shown that Pol IV-dependent 21/22-nt siRNAs are required to silence TEs in microspores. This could be achieved by the non-canonical RdDM pathway involving 21/22-nt siRNAs (Cuerda-Gil and Slotkin, 2016); or, alternatively, by the PTGS pathway. Levels of CHH methylation are low in meiocytes, but increase in microspores and in the vegetative cell of pollen (Walker et al., 2018). Nevertheless, CHH methylation in microspores is very low (Calarco et al., 2012), making it more likely that TE silencing in microspores and later on in the vegetative cell is achieved by PTGS, consistent with the high accumulation of 21/22-nt siRNAs in mature pollen.

Recent work from our and other groups revealed that disruption of NRPD1 suppresses the hybridization barrier between plants of different ploidy grades (Martinez et al., 2018; Satyaki and Gehring, 2019). However, while Martinez et al. (2018) did not find a suppressive effect when using mutants in RdDM components such as *RDR2* and *NRPE1,* Satyaki and Gehring (2019) found those mutants to suppress hybrid seed failure. The difference between both studies lies in the use of tetraploid RdDM mutants by Satyaki and Gehring (2019), while RdDM mutants introgressed into *omission of second division 1 (osd1)* were used by Martinez et al. (2018). Loss of *OSD1* suppresses the second meiotic division, leading to unreduced gamete formation (d’Erfurth et al., 2009). Here, we showed that RDR2 is required for easiRNA biogenesis, suggesting that loss of easiRNAs is not sufficient to suppress the triploid block induced by the *osd1* mutation. An important difference between *osd1* and tetraploid plants is the ploidy of the genome at the beginning of the meiosis, which is diploid and tetraploid, respectively. This fact can have a strong impact, since tetraploid plants undergo DNA methylation changes leading to stable epialleles (Mittelsten Scheid et al., 2003). Given that DNA methylation recruits Pol IV (Law et al., 2013; Zhang et al., 2013) and our study points that easiRNAs are generated during meiosis, it is possible that the requirement of RdDM activity for easiRNA formation and ploidy barriers may be different depending of the initial ploidy of the plants. If true, the signal establishing the triploid block depends on Pol IV but only indirectly on RdDM, suggesting both pathways can be separated, as previously proposed in maize endosperm (Erhard Jr. et al., 2013).

In summary, our study in *Capsella* uncovers a functional requirement of Pol IV in microspores, emphasizing that Pol IV-dependent siRNA formation occurs earlier than previously hypothesized (Slotkin et al., 2009). We show that Pol IV is generating the precursors for 21-24-nt siRNAs, which may be a consequence of different DCLs being able to access Pol IV transcripts during meiosis. Our study highlights the relevance of investigating different plant models to gain novel insights into the molecular control of developmental processes

## Methods

### Plant growth and material

Mutants alleles *nrpd1-3* (Salk_128428) and *dcl3-1* (Salk_005512) have been previously described (Pontier et al., 2005; Xie et al., 2004). For all experiments using *Arabidopsis thaliana,* the Col-0 accession was used as wild type, while for *Capsella rubella,* accession *Cr1GR1* was used.

Seeds of *Arabidopsis* and *Capsella* were surface sterilized in 5% commercial bleach and 0.01% Tween-20 for 10 min and washed three times in sterile ddH2O. Seeds were sown on ½ MS-medium (0.43% MS-salts, 0.8% Bacto Agar, 0.19% MES hydrate and 1% Sucrose). After stratification for 2 days at 4°C, plates were transferred to a growth chamber (16 h light/ 8 h dark; 110 µmol/s/m2; 21°C; 70% humidity). After 10 days, seedlings were transferred to soil and grown in a growth chamber (16 h light / 8 h dark; 110 µmol/s/m2; 21°C; 70% humidity). *Capsella* plants were grown in the growth chamber at the same light-dark cycles, but at 18 °C and 60% humidity.

### Generation of plasmids and transgenic plants

The web tool CRISPR-P (http://cbi.hzau.edu.cn/cgi-bin/CRISPR) was used to design the sgRNAs for knocking out *Capsella NRPD1* (Carubv10019657m) (Lei et al., 2014). Sequence information for the primers containing the two sgRNA sequences are listed in Supplemental table 1. They were used for amplifying the fragment including Target1- sgRNA-scaffold-U6-terminator-U6-29promoter-Target2 using plasmid DT1T2-PCR as template (Wang et al., 2015). The amplified fragment was digested with *Bsa*l and inserted into pHEE401E containing an egg cell specific promoter driven Cas9 cassette as previous described (Wang et al., 2015).

The pHEE401E-*NRPD1*-T1T2 construct was transformed into *Agrobacterium tumefaciens* (GV3101) and bacteria containing the plasmid were used to transform *Capsella rubella* accession *Cr1GR1* by floral dip (Clough and Bent, 1998). The genomic sequence of Arabidopsis *NRPD1* with the stop codon was amplified from Col genomic DNA and cloned into pDONR221 (Invitrogen) and after being confirmed by sequencing it was inserted in pB7WG2 in which the CaMV35S promoter was replaced by the 1.6-kb promoter sequence of *RPS5A* (Weijers et al., 2001).

### Microscopy

*Capsella* inflorescences were harvested and fixed in 3:1 (Ethanol : acetic acid) solution. Pollen were manually dissected from stage 12 and 13 anthers, and then stained with DAPI (1 µg/ml) as previous described (Brownfield et al., 2015). The slides were observed using a Zeiss Axio Scope.A1 and a Zeiss 7800 confocal microscope.

To generate sections, *Capsella* inflorescences were harvested and fixed in FAA solution (50% Ethanol, 5% acetic acid, 4% formaldehyde) and embedded using the Leica Historesin Embedding Kit (702218500). Three-micrometer sections were prepared using a HM 355 S microtome (Microm) with glass knives. Sections were stained with 0.1% toluidine blue for 1 min, washed five times with distilled water, air dried and then observed using Zeiss Axio Scope.

### Microspore extraction

The different pollen stages were extracted on a percoll gradient following previously published procedures (Dupl’akova et al., 2016). The purity of each fraction was assessed by Alexander and DAPI staining.

### RNA and small RNA sequencing

RNA of *Arabidopsis* microspores was isolated using the TRlzoL following the manufacturer’s protocol (Thermofischer: cat-15596018). Purified RNA was treated with DNAsel (Thermofischer: cat-EN0521) and then loaded on a 15% TBE-Urea polyacrylamide gel. RNA with a size of 15-27-nt was retrieved and eluted by crushing the gel in PAGE elution buffer (1M Tris pH7.5, 2.5M NaOAc, 0.5M EDTA pH8) followed by an overnight incubation and a new TRizoL extraction.

*Capsella* leaves were ground with liquid nitrogen, and 100 mg of fine powder from each sample was used for RNA isolation. *Capsella* microspores were ground in a pre-cooled mortar with Lysis/Binding Solution of the *mir*Vana™ miRNA isolation kit. Both long RNAs (>200-nt) and short RNAs (<200-nt) were isolated from leaves and microspores according to the manufacturer’s protocol (*mir*Vana™ miRNA Isolation Kit, AM1560). Size selection of sRNAs was performed as described above.

For the RNA seq analysis, total RNA was treated with the Poly(A) mRNA Magnetic Isolation Module kit (NEB #E7490). Libraries were prepared from the resulting mRNA with the NEBNext® Ultra™ II kit (NEB #E7770S). sRNA seq libraries were generated with the NEBNext® Multiplex Small RNA kit (NEB #E7300S). RNA-seq libraries and sRNA-seq libraries were sequenced at the SciLife Laboratory (Uppsala, Sweden) and Novogene (Hongkong, China) on a HiSeqX in 150-bp paired-end mode or a Illumina HiSeq2000 in 50-bp single-end mode, respectively.

### Bisulfite sequencing

Leaves of 6-10 plants of *Capsella* wildtype and *nrpd1* mutants were pooled as one replicate. Genomic DNA was extracted using the MagJET Plant Genomic DNA Kit (K2761). The bisulfite conversion and library preparation were done as previously described (Moreno-Romero et al., 2016). Libraries were sequenced at the SciLife Laboratory (Uppsala, Sweden) on an Illumina HiSeq2000 in 125-bp paired-end mode.

### DNA sequencing

Genomic DNA was isolated from leaves of one *Capsella* wild-type plant and five *pol iv* mutants using the MagJET Plant Genomic DNA Kit (K2761). Libraries were generated using the NEBNext® Ultra™ II DNA Library Prep Kit for Illumina® and sequenced at Novogene (Hongkong, China) on an Illumina HiSeqX in 150-bp paired-end mode.

### Bioinformatic analysis

For sRNA data, adapters were removed from the 50-bp long single-end sRNA reads in each library. The resulting 18-30-bp long reads were mapped to the respective reference genomes using bowtie (-v 0 --best). All reads mapping to chloroplast and mitochondria and to structural noncoding RNAs (tRNAs, snRNAs, rRNAs, or snoRNAs) were removed. Mapped reads from both replicates were pooled together, sorted in two categories (21/22-nt and 24-nt long) and remapped to the same reference masked genome mentioned above using ShortStack (--mismatches 0 --mmap f) (Johnson et al., 2016) in order to improve the localization of sRNAs mapping to multiple genomic locations. We normalized the alignments by converting coverage values to RPM values. TE-siRNAs were defined as siRNAs that overlap with annotated TEs. TEs accumulating 20 or more reads in the merged wild-type libraries were considered as TE producing siRNA loci. TEs losing siRNAs in *nrpd1* were defined as those having less than 5% of reads left in *nrpd1* compared to wild-type samples. To identify genes losing siRNAs in *nrpd1* microspores, we determined siRNA coverage over the genomic loci plus 2kb up-and downstream regions and calculated differences to wild-type microspores using the Bioconductor RankProd Package (Hong et al 2006) (log2 fold change <-1, (percentage of false prediction) pfp<0.05). For RNA analysis, for each replicate, reads were mapped to the *Arabidopsis* and the *Capsella* reference genomes, using TopHat v2.1.0 (Trapnell et al, 2009) in single-end mode. Gene and TE expression was normalized to reads per kilobase per million mapped reads (RPKM) using GFOLD (Feng et al, 2012). For Capsella the C. *rubella* v1.0 annotated genome was used as reference (Slotte et al., 2013, https://phytozome.jgi.doe.gov/pz/portal.html#!info?alias=Org_Crubella), which was also used as reference in all *Capsella* analyses describe herein. For Arabidopsis the TAIR10 annotation was used. Expression level for each condition was calculated using the mean of the expression values in both replicates. Differentially regulated genes and TEs across the two replicates were detected using the rank product method, as implemented in the Bioconductor RankProd Package (Hong et al 2006). For DNA methylation analysis, reads of each pair were split in 50-bp-long fragments and mapped in single-end mode using Bismark (Krueger and Andrews, 2011). Duplicated reads (aligning to the same genomic position) were eliminated and methylation levels for each condition were calculated averaging the replicates.

To estimate the number and identity of sRNA reads mapping to *COPIA95* TEs in *Capsella,* 21/22-nt and 24-nt sRNA reads were first mapped to a consensus reference fasta file for *Arabidopsis* TEs available at Repbase (https://www.girinst.org/repbase/update/index.html) (Jurka et al 2005) using bowtie (-v 2-m 3 --best --strata). Reads mapping to *COPIA95* TEs were remapped to the C. *rubella* reference genome with ShortStack (-mismatches 0-mmap f) (Johnson et al., 2016) and normalized using coverage values of single copy genes.

New TE insertions in *Capsella rubella* were identified using TEPID (Stuart et al., 2016) in pair-end mode based on the sequenced genomes of five *Cr pol iv* mutants and the corresponding wild-type.

### Data availability

The sequencing data generated in this study are available in the Gene Expression Omnibus under accession number GSE129744. Supplemental Table 2 summarizes all sequencing data generated in this study.

## Author Contributions and Acknowledgments

ZW, NB, and CK performed the experimental design. ZW, NB, JY, and FB performed experiments. GM advised on experimental work. RAM contributed experimental data. ZW, NB, JSG, and CK analyzed the data. ZW, NB, JSG, and CK wrote the manuscript. All authors read and commented on the manuscript.

We are grateful to Cecilia Wärdig for technical assistance. Sequencing was performed by the SNP&SEQ Technology Platform, Science for Life Laboratory at Uppsala University, a national infrastructure supported by the Swedish Research Council (VRRFI) and the Knut and Alice Wallenberg Foundation. This research was supported by grants from the Swedish Research Council VR and Formas (to CK), a grant from the Knut and Alice Wallenberg Foundation (to CK), and support from the Göran Gustafsson Foundation for Research in Natural Sciences and Medicine (to CK). Research in the Martienssen laboratory is supported by the US National Institutes of Health (NIH) grant R01 GM067014, and by the Howard Hughes Medical Institute. The authors acknowledge assistance from the Cold Spring Harbor Laboratory Shared Resources, which are funded in part by the Cancer Center Support Grant (5PP30CA045508).

## Supplemental Figure legends

**Supplemental Figure 1.**
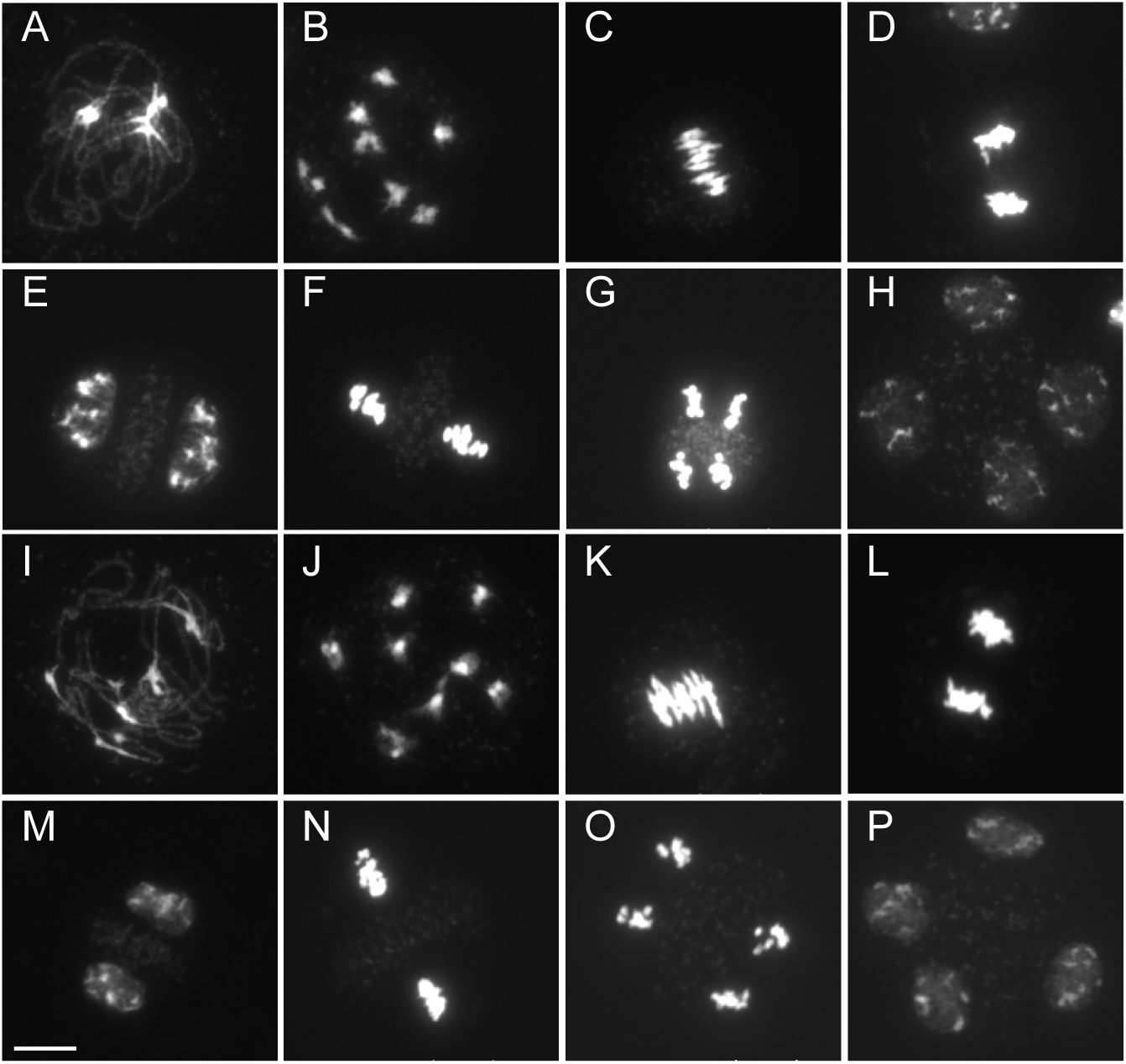
Meiosis is not affected in *Capsella nrpd1.* Supports Figure 2. Meiosis in *Capsella* wild-type (A - H) and *nrpd1* (I - P) plants. A and I, pachytene. B and J, diakinesis. C and K, metaphase I. D and L, telophase I. E and M, prophase II. F and N, metaphase II. G and O, anaphase II. H and P, telophase II. Bar: 5 µm.

**Supplemental Figure 2.**
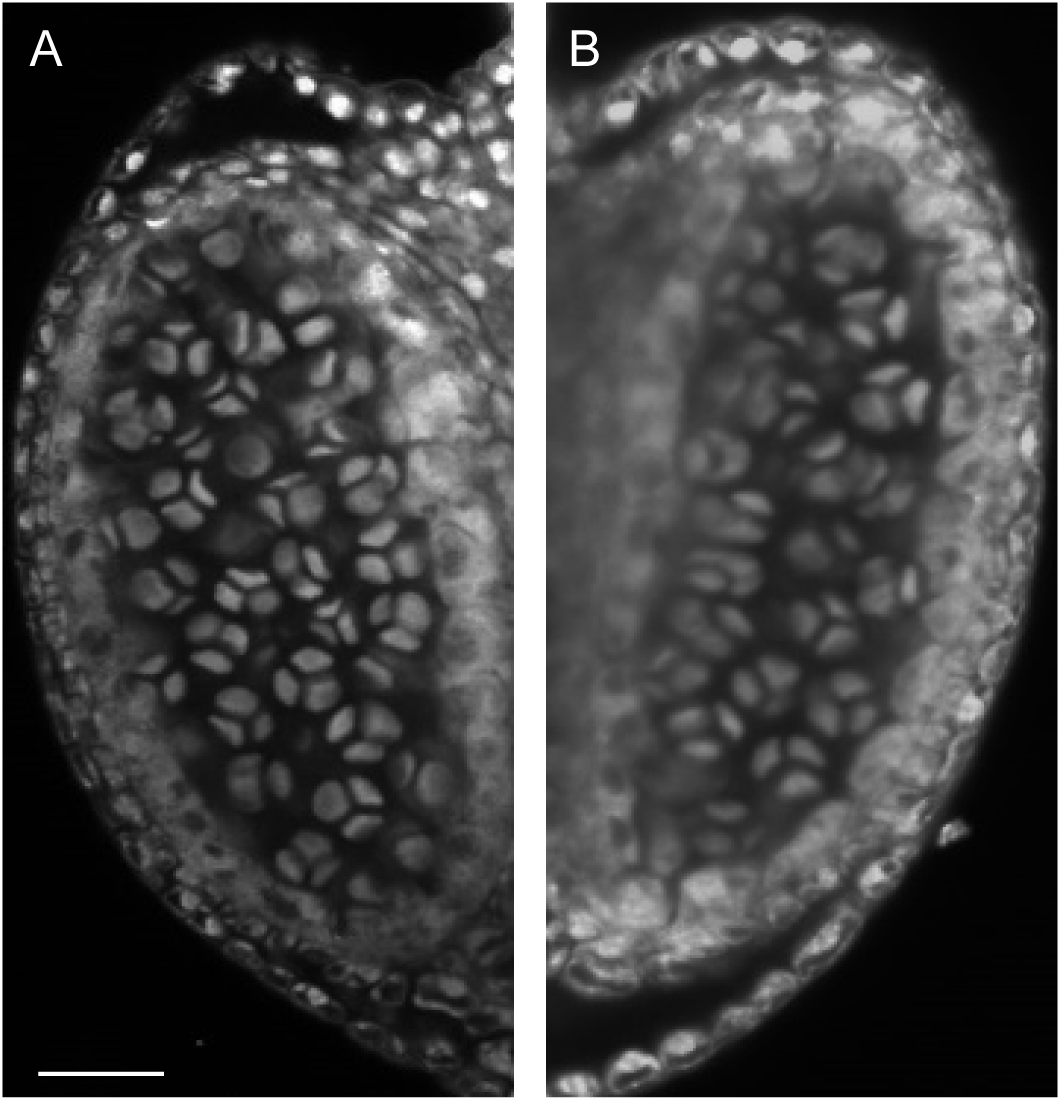
Normal tetrad formation in *Capsella* wild type (A) and *pol iv* (B). Supports Figure 2. Shown are whole mount confocal images. Bar: 20 µm.

**Supplemental Figure 3.**
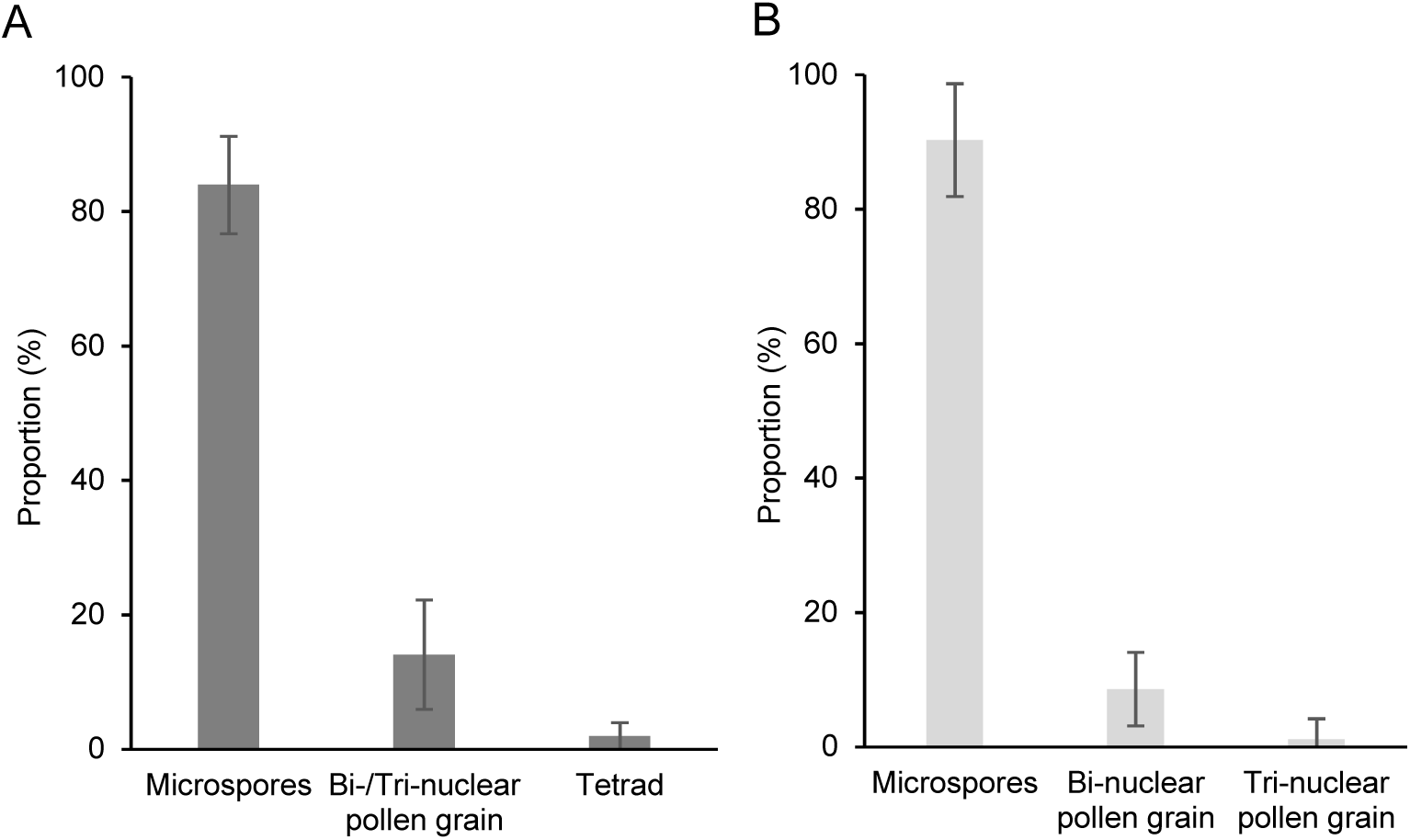
Average purity of *Capsella* and *Arabidopsis* microspore extractions. Supports Figure 4. Microspore extractions of *Capsella* (A) and *Arabidopsis* (B) were tested by DAPI staining and the B2 and B1 fractions were selected as the fractions containing the highest proportion of microspores in *Capsella* and *Arabidopsis,* respectively. Shown is the average percentage of four and eight independent extractions in *Capsella* and *Arabidopsis,* respectively. Error bars show standard deviation.

**Supplemental Figure 4.**
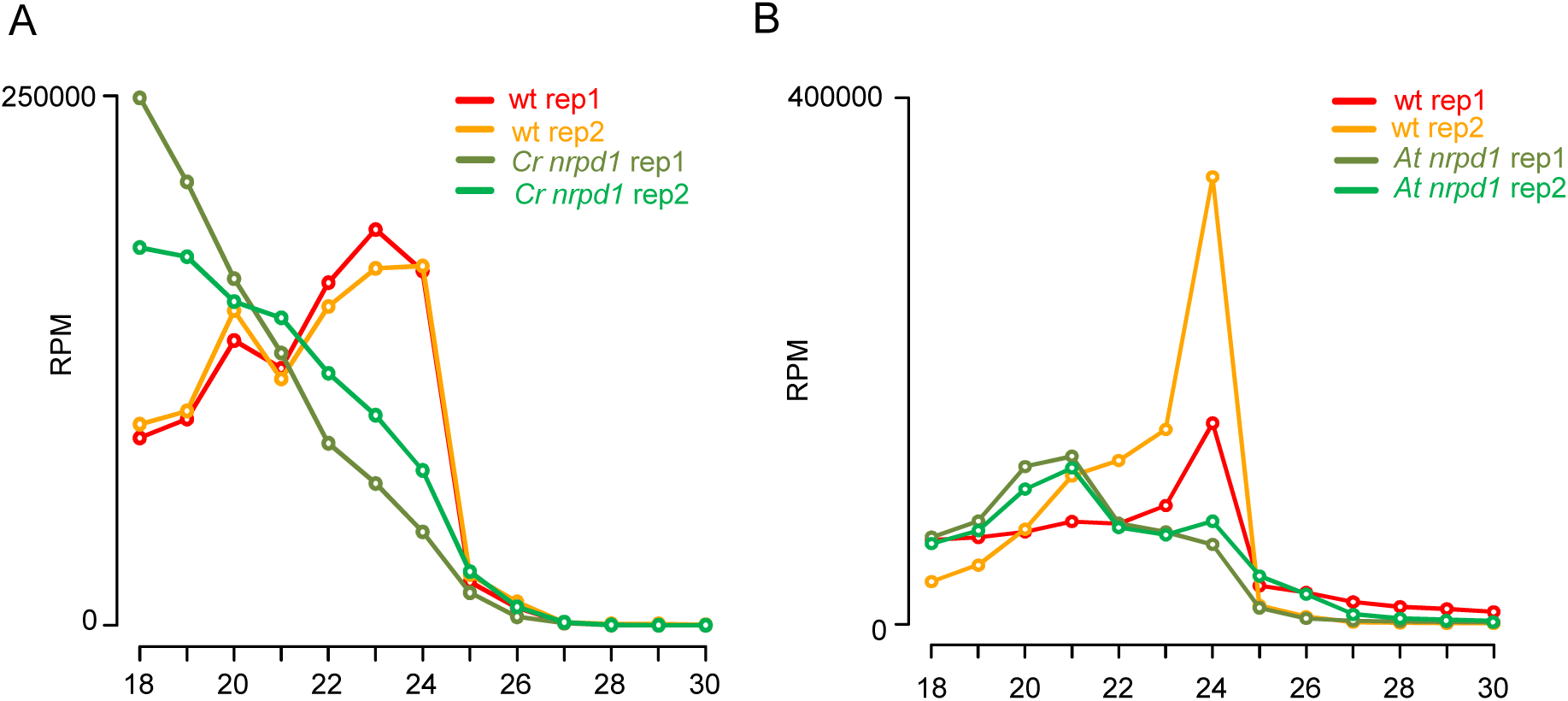
Profile of total sRNAs in *Capsella* (A) and *Arabidopsis* (B) microspores. Supports Figure 4.

**Supplemental Figure 5.**
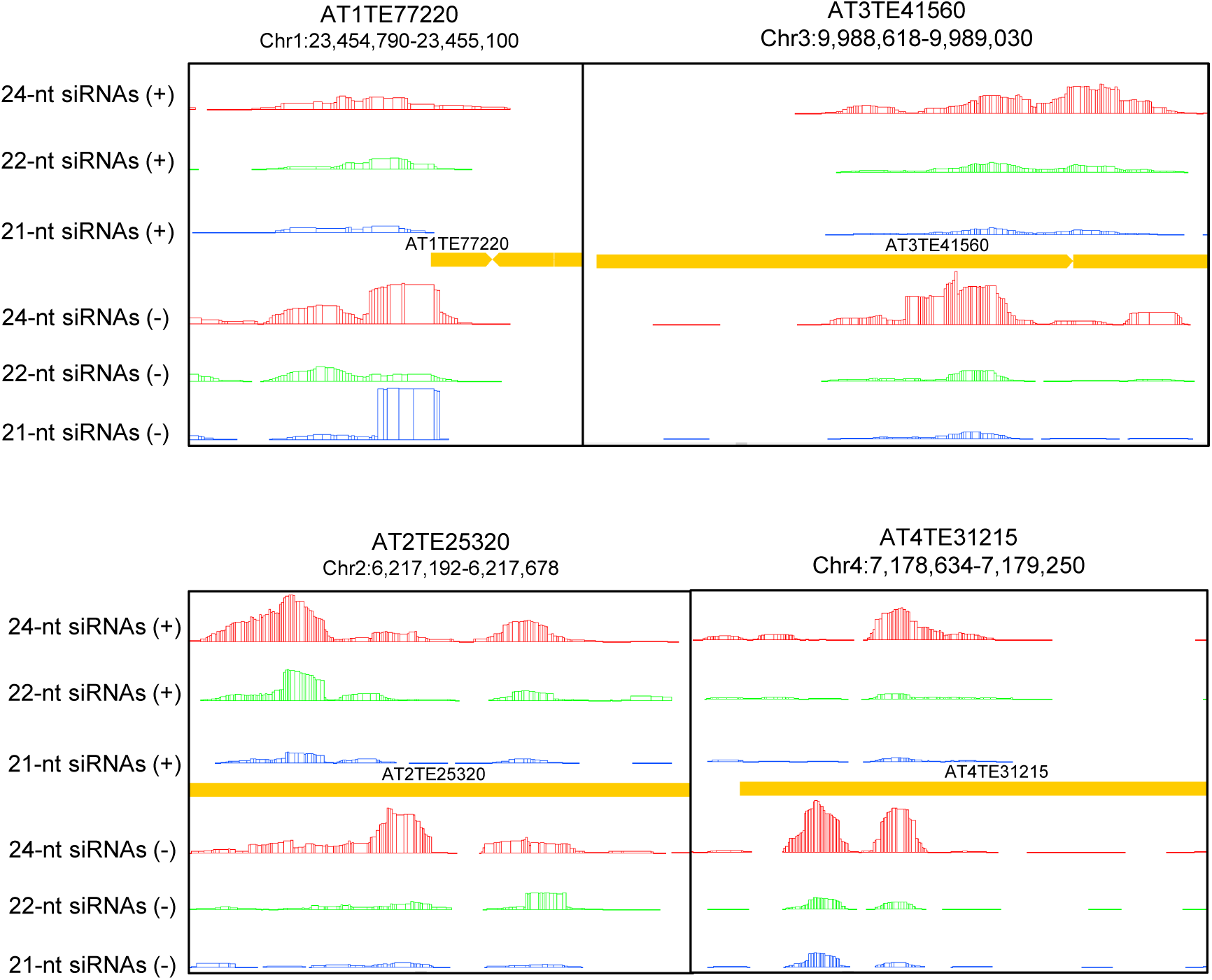
Example of four loci producing Pol IV-dependent siRNAs in *Arabidopsis.* Supports Figure 7. Bars represent normalized reads. The color indicates the length of the analyzed reads: red 24-nt, blue 22-nt, and green 21-nt. The DNA strand is indicated by the(+) or (-). TE sequences are represented in yellow.

**Supplemental Figure 6.**
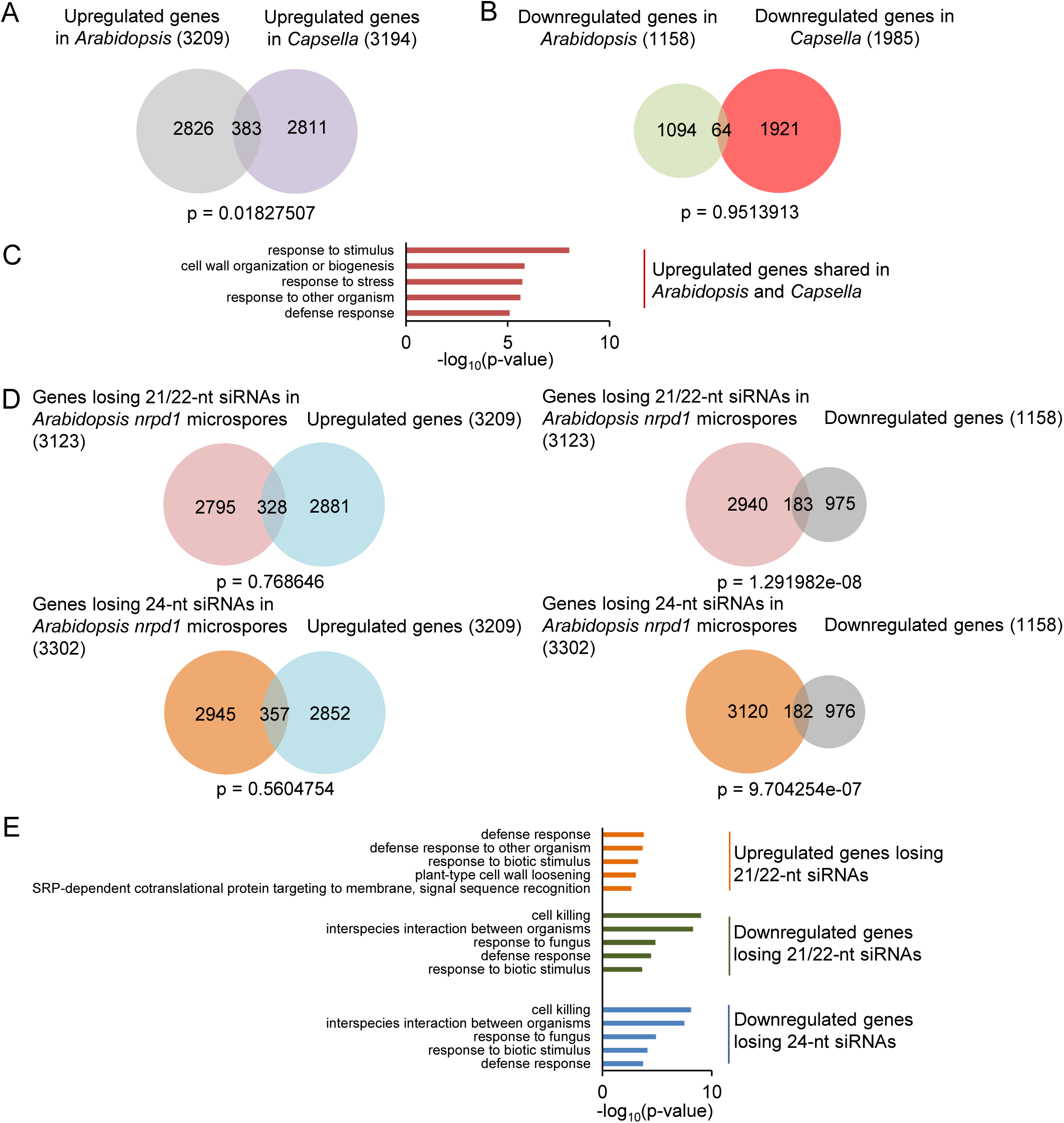
Deregulated genes in *Arabidopsis nrpd1* mutant microspores. Supports Figure 8. (A) Venn diagram showing overlap of upregulated genes in *nrpd1* microspores of *Capsella* and *Arabidopsis.* Significance was determined by a hypergeometric test. (B) Venn diagram showing overlap of downregulated genes in *nrpd1* microspores of *Capsella* and *Arabidopsis.* Significance was determined by a hypergeometric test. (C) Enriched gene ontologies (GOs) for biological processes of upregulated genes shared in *Arabidopsis* and *Capsella nrpd1* microspores. Top 5 GOs are shown. (D) Venn diagrams showing overlap of deregulated genes (|log_2_ fold change|> 1, p < 0.05) in *nrpd1* microspores of *Arabidopsis* and genes losing 21/22-nt and 24-nt siRNAs at 2kb up-and downstream and gene body (log_2_ fold change < −1, p < 0.05) in *Arabidopsis nrpd1* microspores. Significance was determined by a hypergeometric test.

**Supplemental Figure 7.**
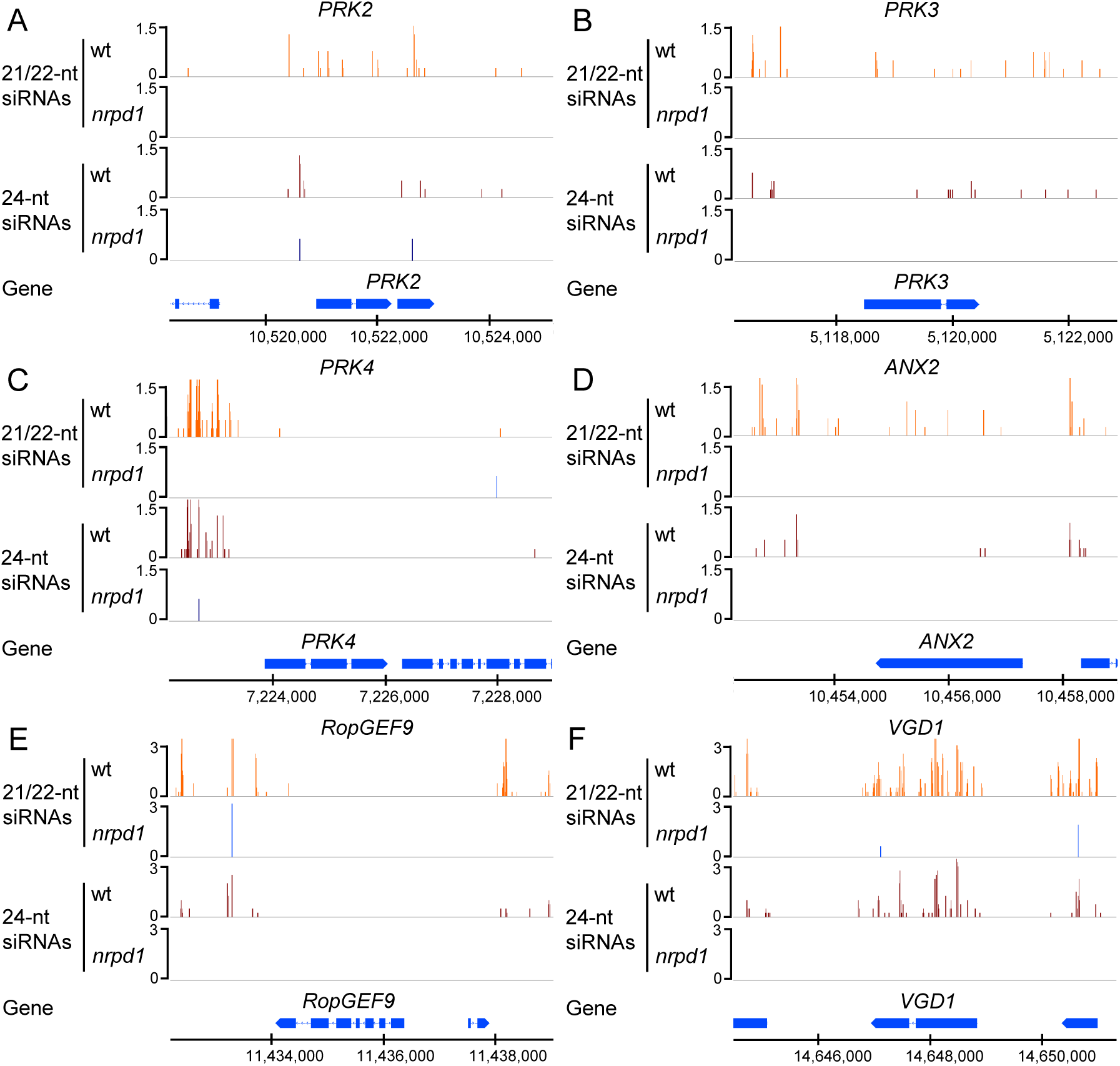
Representative pollen developmental genes accumulating 21/22-nt and 24-nt siRNAs in *Capsella* microspores.

## Supplemental datasets

**Supplemental dataset 1. *COPIA95* elements accumulating Pol IV-dependent siRNAs in *Capsella* and *Arabidopsis*.**

**Supplemental dataset 2. Up-and downregulated genes in *Arabidopsis* and *Capsella* microspores.**

**Supplemental dataset 3. Up-and downregulated genes in *Capsella* microspores overlapping with regions losing 21/22-or 24-nt siRNAs in *Cr nrpd1* microspores. Supplemental dataset 4. Up-and downregulated genes in *Arabidopsis* microspores overlapping with regions losing 21/22-or 24-nt siRNAs in *At nrpd1* microspores.**

---

**Supplemental table 1.**
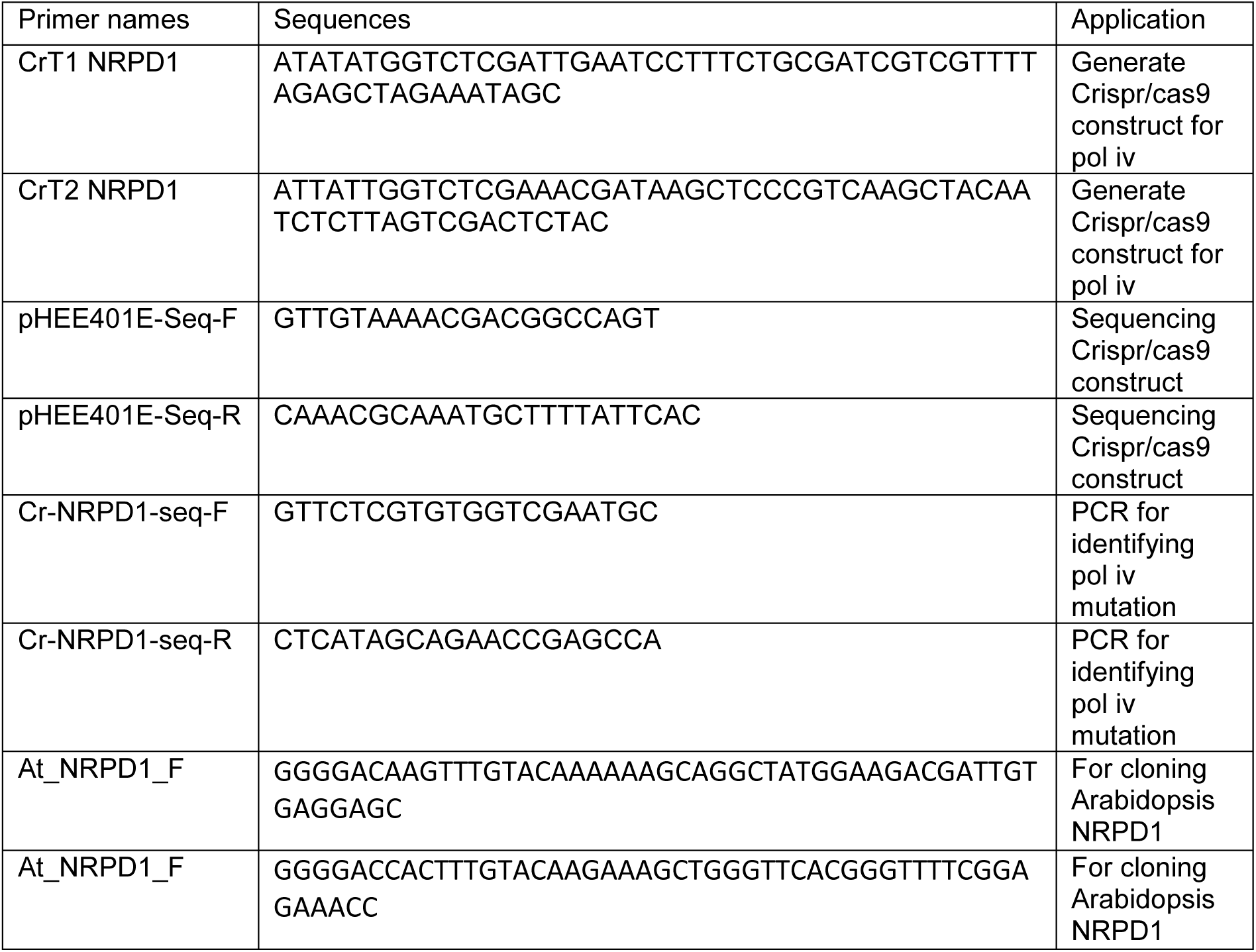
Primer list.

**Supplemental table 2.**
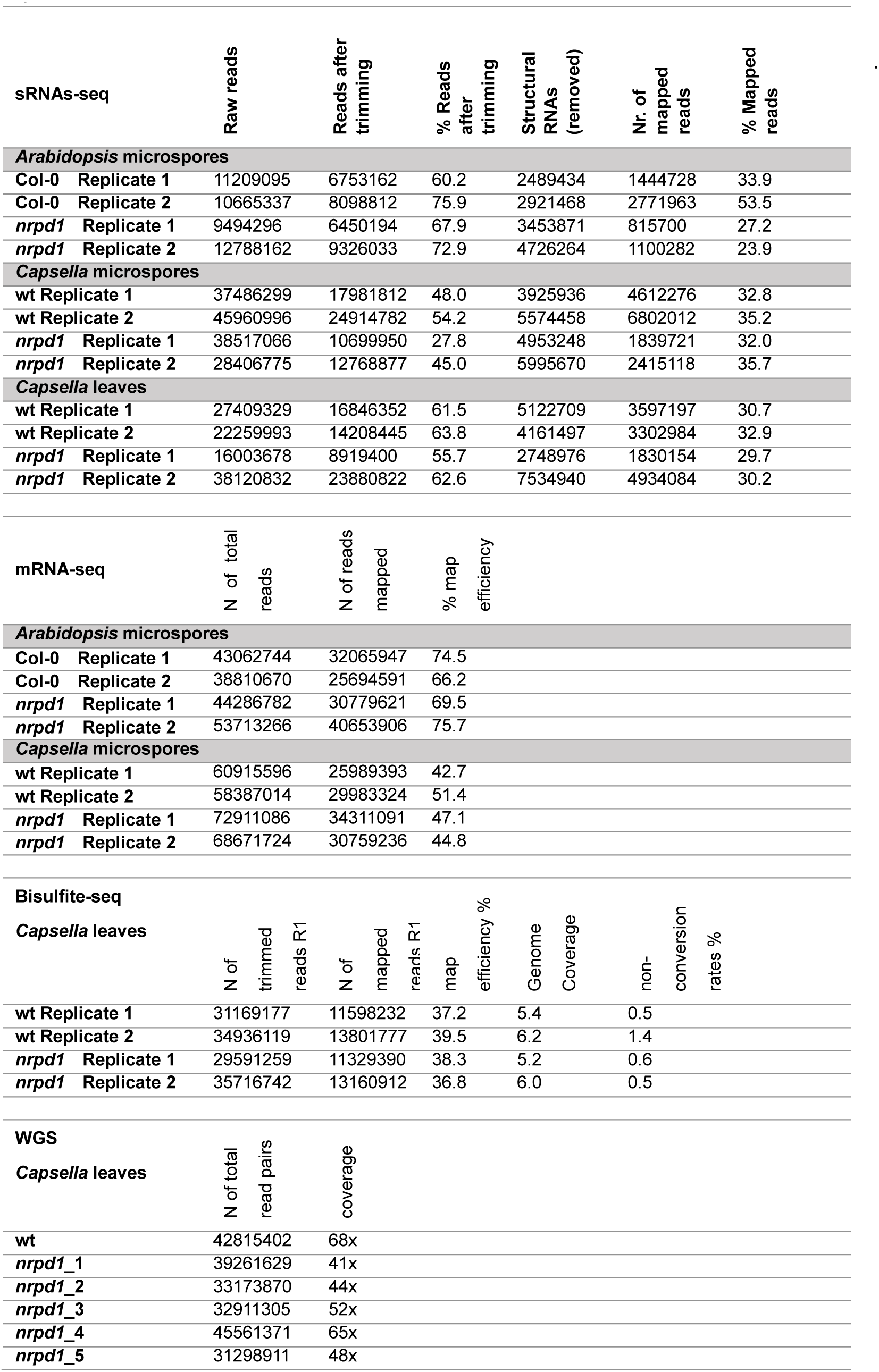
Quality of sequencing samples. Table shows details of the sequenced samples generated in this study. Replicates are biological replicates.

